# The CD19-4-1BBL antibody fusion protein unleashes the immune system against high-risk chronic lymphocytic leukemia

**DOI:** 10.64898/2026.07.09.737515

**Authors:** Priyanka Khare, Ronghua Zhang, Cristina Ivan, Sarah Schneider, Priyanka Banerjee, Navid Sobhani, Harry Polasek, Vanessa Behrana Jensen, Karen Clise-Dwyer, William Wierda, Maria Teresa Sabrina Bertilaccio

## Abstract

CD19-4-1BBL is a bispecific antibody fusion protein that targets CD19 and costimulates 4-1BB on T cells and other immune cells. Its antitumor activity has been reported in B-cell non-Hodgkin lymphoma with emphasis on its T-cell mediated cytotoxic activity. Its effect on other 4-1BB expressing immune cells is unexplored. Here, we investigated the molecular mechanisms and the antileukemic effect of CD19-4-1BBL in chronic lymphocytic leukemia (CLL), a B-cell malignancy profoundly marked by the immunosuppressive activity of myeloid-derived suppressor cells, tumor-associated macrophages and CD4^+^ regulatory T cells. We demonstrated that CD19-4-1BBL simultaneously mitigates the immunosuppressive phenotype and transcriptome machinery of these cells and promotes antitumor CD8^+^ T-cell immunity. Finally, in a preclinical, patient-derived xenograft model of CLL, we observed a favourable survival impact, especially in mice transplanted with immune cells from patients with high-risk/progressive leukemia. Our findings provide evidence that the CD19-4-1BBL treatment is a multifaceted, immune-based strategy that should be clinically explored in patients with chronic lymphocytic leukemia.

**KEY POINTS:**

- CD19-4-1BBL sharpens the myeloid cell transcriptome and stimulates diverse memory CD8^+^ T cell clonotypic responses.
- CD19-4-1BBL costimulation can be therapeutically exploited in high-risk chronic lymphocytic leukemia.

## INTRODUCTION

CLL cells strongly depend on survival signals provided by the non-malignant immune microenvironment. T cells in patients with CLL are abnormal, with profound functional defects and features of exhaustion ^1, 2^. Immunosuppressive regulatory CD4^+^ FoxP3^+^ T cells (T_REGS_) have been identified in patients with CLL and, as in solid tumors, suppress the antitumor response of cytotoxic T cells ^3^. Within the myeloid cell compartment, monocytes, tumor associated macrophages (TAMs) and myeloid derived suppressor cells (MDSCs) have crucial immunosuppressive functions and help leukemic cells to growth and proliferate ^4,5,6^. We recently demonstrated that CLL patient-derived monocytes and MDSCs differentiate into TAMs ^7^. Altogether, T_REG_ cells, MDSCs, TAMs and their progenitors nurture the immunosuppressive microenvironment.

We have selected the antibody fusion protein CD19-4-1BBL with the purpose to target the immunosuppressive cells of the immune microenvironment and repair dysfunctional T cells. CD19-4-1BBL is composed of split trimeric 4-1BB ligands and a CD19-targeting portion, fused to a silent Fc part.^8^ In the presence of T-cell receptor (TCR) signaling, this bispecific molecule provides potent, tumor-antigen-mediated T-cell costimulation.

4-1BB is expressed by several cell types in the immune system^9^ and promotes the cytotoxic function of T cells and natural killer (NK cells).^10,11^ In mice, 4-1BB-4-1BBL interactions are also involved in myeloid-lineage cellular development and differentiation.^12^

4-1BB is a member of the tumor necrosis factor (TNF)-receptor superfamily. Upon 4-1BB activation through crosslinking with an agonist, TNF receptor-associated factors (TRAFs) are recruited to the cytoplasmic domain of 4-1BB and initiate the construction of the 4-1BB signalosome.^13^ The architecture of the 4-1BB-TRAF signaling complexes in each cell type depends on the activation state of the cell and on the relative expression level of TRAFs.^13^ Many proteins are recruited to 4-1BB-TRAF complexes including ligases, proteases, kinases, and modulatory proteins. The 4-1BB signalosome formation can result either in the activation of nuclear factor-kappa B (NF-κB) and extracellular signal–related kinase (ERK), or the c-JUN N-terminal kinase (JNK) and p38MAP kinase cascades. Overall, 4-1BB-4-1BBL costimulation supports the survival of T cells.^14^

Unlike memory T cells, the role of 4-1BB signalosome in human T_REG_ and myeloid cells is undefined. Besides its involvement in antitumor immunity, the 4-1BB-4-1BBL signaling has been investigated in antiviral immunity. 4-1BBL or 4-1BB deficiency hampers CD8^+^ T cell response during acute infections with VSV, LSMV and some influenza strains.^14^

Prompted by these observations, we investigated the molecular mechanisms whereby CD19-targeted 4-1BB costimulation in CLL affects immune cells of the adaptive and innate immunity. We focused on CD8^+^ memory T cells, CD4^+^ T_REGS,_ monocytes/macrophages and monocytic-MDSCs (M-MDSCs). We confirmed the activation and costimulation of leukemia-specific memory T cells at the transcriptional level. Upon CD19-4-1BBL, we observed a mitigation of the immunosuppressive activity of CD4^+^ T_REG_ cells, TAMs and M-MDSCs at transcriptional and protein level. In combination with the glycoengineered type II CD20 monoclonal antibody (mAb) obinutuzumab,^15^ the therapeutic CD19-4-1BBL bispecific immunomodulator favorably impacted the survival of or cured xenotransplanted leukemic mice adoptively transferred with immune cells derived from patients with high-risk CLL. Our findings critically support the therapeutic use of CD19-4-1BBL in patients with high-risk CLL.

## MATERIALS AND METHODS

### Cells and reagents

Human primary samples were obtained from patients with CLL (all Rai stages) referred to the Leukemia Department at The University of Texas MD Anderson Cancer Center with the approval of MD Anderson’s Institutional Review Board (protocol LAB04-0678) and, in accordance with the Declaration of Helsinki. Written informed consent was obtained from the donors. The clinical and biological features of the patients analyzed are described in Supplementary Tables S1, S2. All the patients were either untreated or off therapy for at least 8 months before the beginning of the study. The MEC1 cell line^16^ was obtained from Deutsche Sammlung von Mikroorganismen und Zellkulturen (DMSZ, Braunschweig, Germany) and is described in the Supplementary Methods. Anti-human CD19-4-1BBL^8^ bispecific fusion protein and anti-human CD20 obinutuzumab^15^ were provided by Roche Innovation Center (Zurich, Switzerland) or purchased at MedChemExpress (Monmouth Junction, NJ, USA).

### In vitro cultures

Fresh peripheral blood mononuclear cells (PBMCs) from untreated CLL patients were seeded at 3 x 10^6^ cells/mL in culture medium and treated with either CD19-4-1BBL (10 g/mL, Roche), or CD20 obinutuzumab (10 g/mL), or both. Fluorescence-activated cell sorting was performed after 22h of treatment (described below and in the Supplementary Methods). The in vitro cultures for TCR sequencing studies and quantitative flow cytometry-based cell-depletion assays are described in the Supplementary Methods and Supplementary Tables S3 and S4.

### Human cell sorting

Fluorescence-activated cell sorting of live human myeloid cells and of live human lymphoid cells was performed using BD FACSAria Fusion and BD Influx instruments (BD Biosciences, Franklin Lakes, NJ). Methodologies and antibodies are described in the Supplementary Methods.

### Conventional and spectral flow cytometry

Conventional flow cytometry phenotype analysis of live human myeloid cells and of live human lymphoid cells was performed using an LSRFortessa X-20 instrument (BD Biosciences).

Spectral flow cytometry analysis was performed using a Cytek Aurora instrument (Cytek Biosciences, Fremont, CA). Flow cytometry data were analyzed with FCS Express 6 or FCS Express 7 Flow Cytometry software (De Novo Software, Pasadena, CA). Methodologies and antibodies are described in the Supplementary Methods and Tables S3-S5.

### Gene expression profiling and RNA TCR sequencing analysis

After sorting of myeloid and lymphoid cells, RNA extraction was performed using an RNeasy Mini Kit (QIAGEN, Hilden, Germany). Microarray analysis (Human Clariom D Pico Assay) was performed at the MD Anderson’s Sequencing and Non-Coding RNA Program. TCR sequencing (SMARTer Human TCR a/b Profiling Kit v2) was performed at the MD Anderson’s Cancer Genomics Laboratory. For detailed methodologies and analysis see Supplementary Methods.

### Mice

MISTRG mice were kindly provided by Regeneron (Tarrytown, NY). MISTRG mice are on the Rag2-/-gc-/-background and carry genes encoding human macrophage colony-stimulating factor (MCSF), human interleukin-3 (IL-3) and granulocyte-macrophage colony-stimulating factor (GM-CSF), human thrombopoietin (TPO) and a bacterial artificial chromosome (BAC)-transgene encoding human SIRP-.^17^ Mice were housed and bred in a specific pathogen-free animal facility at MD Anderson Cancer Center. They were treated in accordance with the approval of the Institutional Animal Care and Use Committee of MD Anderson Cancer Center (protocol 00001627-RN02), and the study was conducted in accordance with the Animal Welfare Act, the Guide for the Care and Use of Laboratory Animals, and the Public Health Service (PHS) Policy. For detailed information on xenograft studies and murine cell preparations see Supplementary Methods.

### Statistical analysis and Graphical Representation

The statistical analysis and graphical representation of the data was performed using the GraphPad Prism 9.0 and R version 4.4.0 software. For detailed description of the methodology, including microarray and TCR sequencing processing, see Supplementary Methods.

## RESULTS

### CD19-4-1BBL TRIGGERS THE 4-1BB SIGNALOSOME OF MEMORY T CELLS FROM PATIENTS WITH CLL

To evaluate the effect of CD19-4-1BBL on immune cells from patients with CLL, a broad transcriptome-gene- and exon-level analysis of coding and ncRNA isoforms was performed using the Affimetrix Clariom D assay on CD14^+^ monocyte subsets, CD14^+^ HLA-DR^low/-^ M-MDSCs, effector (T_EM_) and central memory (T_CM_) CD8^+^ T cells, and regulatory CD4^+^ T_REG_ isolated from patients with CLL (**Table S1-S2**, **Figure 1A**). Monocytes subsets were identified as CD14^+^ CD16^++^ nonclassical (NC), CD14^++^ CD16^+^ intermediate (I), and CD14^++^ CD16^-^ classical (C) monocytes; CD8^+^ T_EM_ and T_CM_ cells were identified based on the differential surface expression of CD45RO, CD45RA, and CD62L.^18^ These immune cells from patients with CLL express 4-1BB (**Figure S1A**). Principal-component analysis (PCA) segregated the transcriptomic profile of T lymphocytes from that of myeloid cells (**Figure 1B**). A number of transcripts were modulated not only on T cells but also on myeloid cells (**Table S7**). Common transcripts were found up- or down-modulated on immune cells with immunosuppressive function either from the lymphoid or myeloid lineage, including M-MDSCs, T_REGS_, and I monocytes (**Figure 1C, Table S8).** Gene ontology (GO, **Table S9**) and Pathway (**Table S10**) analyses were performed for the genes significantly modulated upon CD19-4-1BBL treatment in each cell type compared to the untreated control by using the web tool Enrichr.

**Fig. 1.**
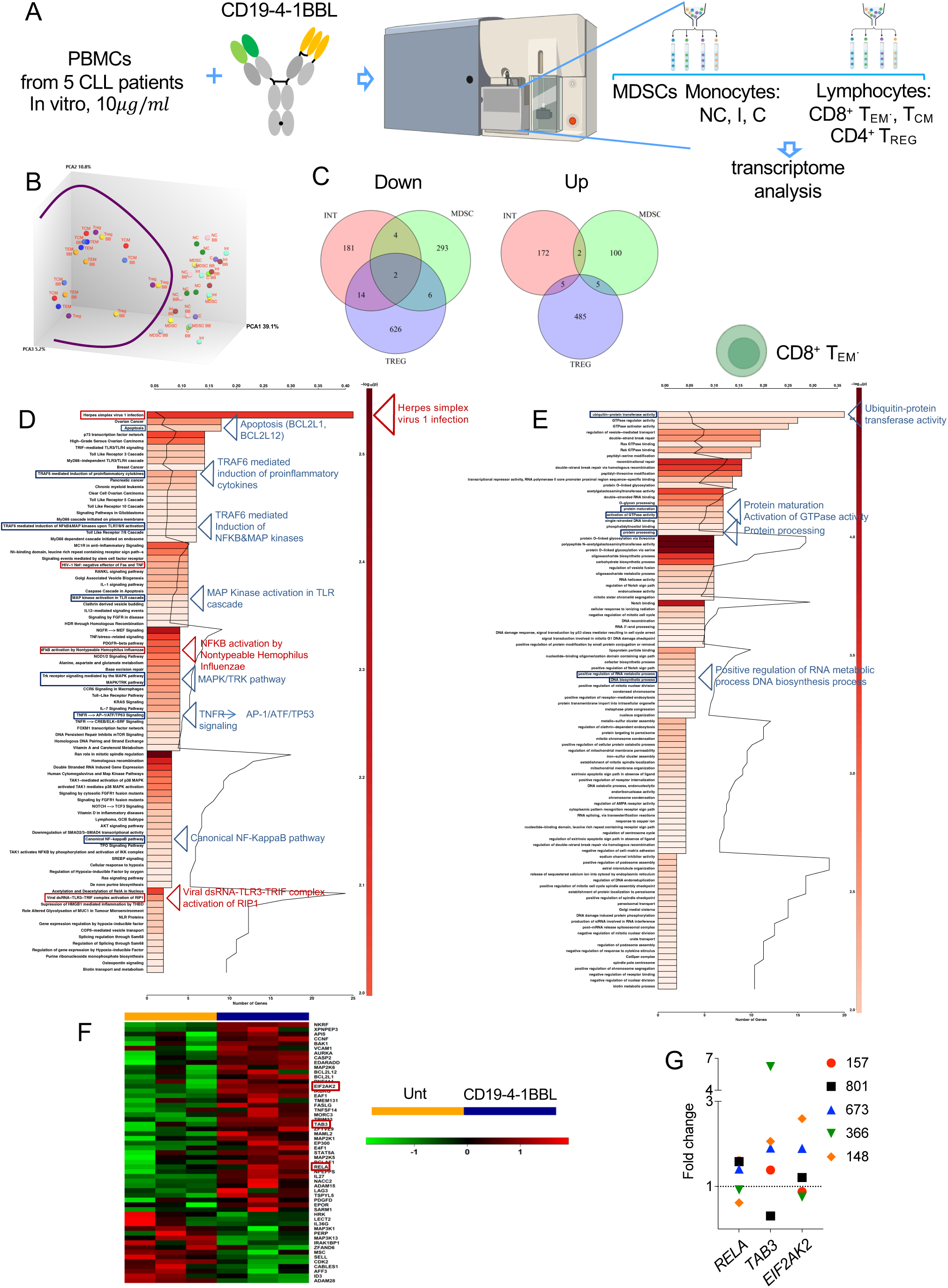
Transcriptome analysis of patient-derived immune cells exposed to CD19-4-1BBL. (A) Fresh peripheral blood mononuclear cells (PBMCs) from patients with chronic lymphocytic leukemia (CLL; n=4; Table S1) were incubated with CD19-4-1BBL (10µ/mL) for 22 h. RNA was isolated from fluorescent-activated cell sorted (FACS) CD19^+^ B cells, hCD8^+^ CD45RA^-^CD45RO^+^CD62L^-^ effector memory T cells (T_EM_), hCD8^+^ CD45RA^-^CD45RO^+^CD62L^+^ central memory T cells (T_CM)_, CD4^+^CD25^+^CD127^low/-^ regulatory T cells (T_REGS_), CD14^+^ CD16^++^ nonclassical (NC) monocytes, CD14^++^ CD16^+^ intermediate (I) monocytes and CD14^++^ CD16^-^ classical (C) monocytes and CD14^+^HLADR^low/-^ monocytic myeloid-derived suppressor cells (M-MDSCs). A human Clariom D Pico assay was used to perform a broad transcriptome gene- and exon-level analysis of coding RNA. (B) Principal-component analysis (PCA) plot of genes expressed on lymphoid and myeloid cells treated with CD19-4-1BBL or left untreated. Each color on the panel identifies a cell type (total samples = 44). (C-D) Venn diagrams of commonly downregulated and upregulated transcripts in I monocytes, M-MDSCs and CD4^+^ T_REGS_ are shown. (D) Gene ontology (GO) and (E) pathway analysises of differentially expressed transcripts between CD19-41BBL-treated and untreated CD8^+^ T_EM_ are shown. (F) A supervised heatmap for chosen, differentially expressed transcripts for CD8^+^ T_EM_ generated in R using the heatmap.2 function of the gplots library is shown. (G) Relative mRNA expression of *RELA, TAB3* and *EIF2AK2* in human primary CD8^+^ T_EM_ separated from PBMCs by FACS (n = 5, patients 157, 801, 673, 366, 148, Supplementary Table S1) and plated in 6-well plates alone or with 10 μg/mL of CD19-4-1BBL for 22 h. Three technical replicates were analyzed for each sample. Data were normalized to β-actin expression. Gene expression was determined by calculating the difference (ΔCt) between the threshold cycle (Ct) of each gene and that of the reference gene and was expressed as the mean of 3 replicates ± the standard error of the mean (SEM). Then the relative quantification values were calculated as the fold change expression of the gene of interest over its expression in the selected cell-type reference sample, i.e., the untreated sample (considered as the calibrator sample), using the formula 2 ^-ΔΔCt^. Finally, the treated sample’s relative mRNA expression was normalized to the vehicle. Samples with undetermined Ct values were not included.

As expected, we found that CD8^+^ T_EMS_ upregulated transcripts and pathways involved in the 4-1BB signalosome cascade, including TRAF6-mediated induction of proinflammatory cytokines, NF-κB and MAP kinases, TNFR signaling, canonical NF-κB pathway (**Figure 1D**). These findings were confirmed by the GO analysis reporting a number of enriched terms known to be induced by the 4-1BB-4-1BBL costimulation, including ubiquitin-protein transferase activity, protein maturation and processing, activation of GTPase activity, positive regulation of RNA metabolic process and DNA biosynthesis process (**Figure 1E**). Of interest, genes involved in infections and antiviral immunity were upregulated (**Figure 1D**). Among the differentially expressed genes, we selected a limited number of transcripts, somehow involved in the 4-1BB-4-1BBL costimulation or with relevant function for the specific cell type, and these were validated by RT-PCR. Supervised heatmaps for selected genes were generated by R language (**Figure 1F-G, Table S11**). The pathway analysis of CD8^+^ T_CM_ confirmed the activation of the 4-1BB signalosome through the IKK/NF-κB signaling pathway (**Figure S1B**). The GO analysis highlighted the enrichment of the 4-1BB signalosome-related JNK cascade (**Figure S1C**).

### TCR SEQUENCING OF CD8^+^ MEMORY T CELLS REVEALS CLONOTYPE INDUCTION UPON CD19-4-1BBL STIMULATION

Given the transcriptional changes observed on memory T cells, we asked whether CD19-4-1BBL induced expansions of specific clones of CD8^+^ T_EM_ cells. Therefore, we performed TCRα/β RNA profiling of CD8^+^ T_EM_ separated from the PBMCs of patients with CLL to identify clonotypes induced by the combination of CD19-4-1BBL and αCD20 obinutuzumab (**Figure 2A**). αCD20 was used to induce CLL cell death and provide Antigen (Ag) stimulation. We found that treated T_EM_ cells harbored increased counts of clonotypes compared with the untreated samples (**Figures 2B-C, Tables S12-S13**). Additionally, we observed an increase in the *TRA* and *TRB* CDR3 length distributions (Figure **S2A, B)**, indicating enhanced diversity and broader range of Ag-recognition capabilities of CD8^+^ T_EM_ cells treated with CD19-4-1BBL. We next analyzed the *TRA/TRB* genes’ CDR3 sequences and their related tumor-associated antigens (**Tables S14-S15**) mainly through the VDJ database (vdjbase.org) and as a result, we identified CLL-associated Ags, including BST2 and SF3B1. SF3B1 is a crucial component of the splicing machinery. Mutations in SFRB1 have been identified in patients with CLL at high frequencies and are associated with aggressive disease and shorter survival ^19,20^. BST2 is a lipid-raft-associated protein and has been reported to be over-expressed on CLL cells ^21^. Of note, together with CLL-specific Ags, we also identified solid tumor-associated Ags (TAA), including MLANA and PMEL (**Tables S14, S15**). We also found a number of virus-specific Ags related to EBV, CMV, HCV, HIV, SARS-Cov-2 (**Tables S16-S21,** *TRB*), demonstrating that CD19-4-1BBL and αCD20 combination potentiates wide antitumor and antiviral T-cell specific immunity.

**Fig. 2.**
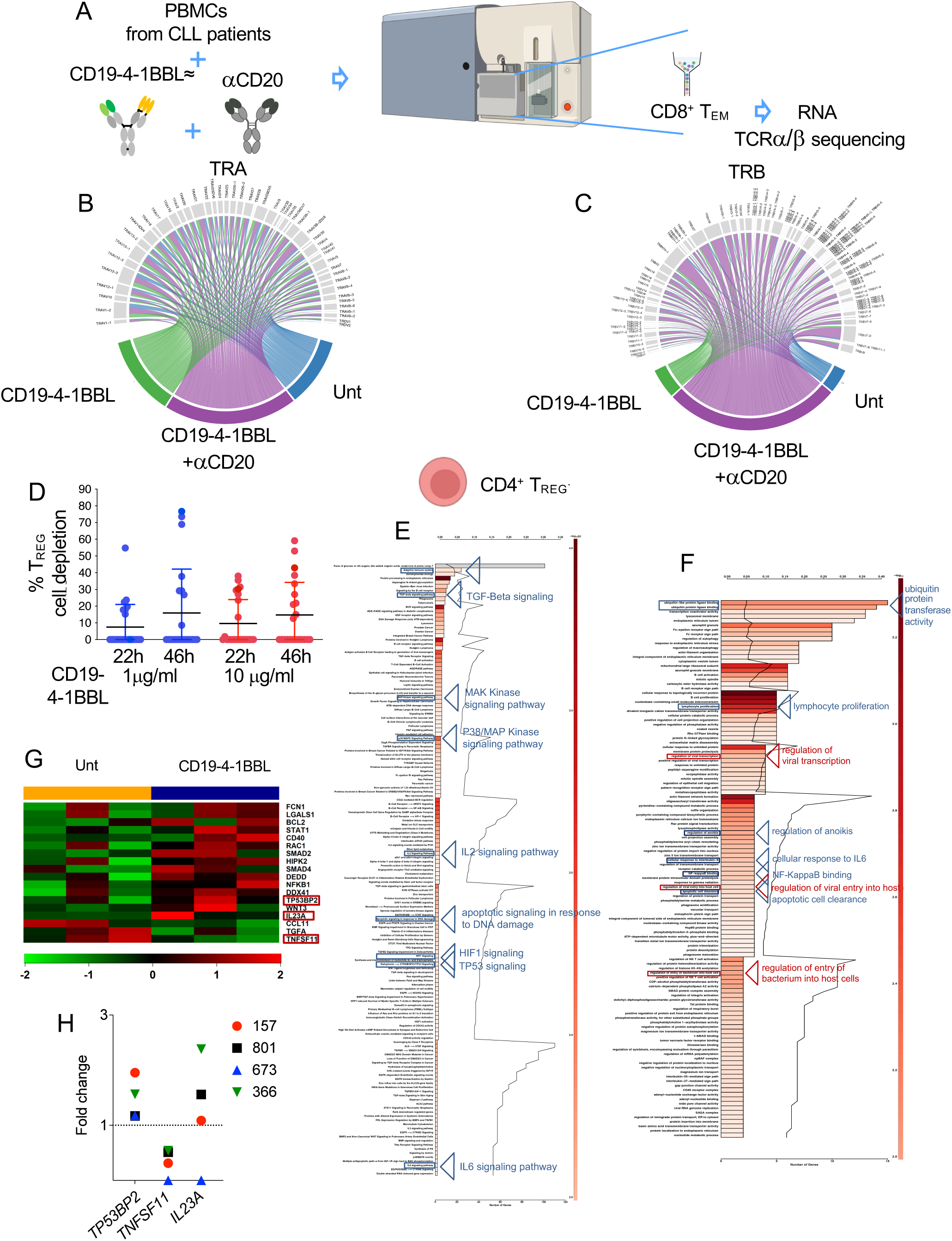
Costimulatory activity of CD19-4-1BBL in patient-derived effector memory CD8+ T cells and CD4+ TREG cells. (A) Fresh PBMCs from patients with CLL (n = 3, Table S1) and age-matched healthy donor controls (n = 3) were incubated with CD19-4-1BBL (10µ/mL) or CD19-4-1BBL plus CD20 obinutuzumab (10 µg/mL) or left untreated for 48 h. RNA was isolated from FACS sorted hCD8^+^ CD45RA^-^CD45RO^+^CD62L^-^ T_EM_ and the Takara SMARTer Human TCR a/b Profiling Kit v2 was performed. (B-C) The circos plots show the clonotype distributions and rearrangements of the TCR and TCR chains, respectively. Complete information including abbreviations and TRA and TRB rearrangements is shown in Tables S8-S9. (D) hCD4^+^CD25^+^Foxp3^+^ T_REG_ cell depletion (n = 25, Supplementary Table S2) after 22 h and 46 h of treatment with 1µg/mL and 10µg/mL of CD19-4-1BBL was analyzed using multi-color flow cytometry and calculated by the formula: the following formula: 100 - % remaining cells, where % remaining cells = (absolute number in treated samples/absolute number in untreated samples) × 100. Details are described in the Supplementary Methods. (E) GO and (F) pathway analyses of differentially expressed transcripts between CD19-41BBL treated and untreated CD4^+^ T_REG_ are shown. (G) A supervised heatmap for chosen, differentially expressed transcripts on CD4^+^ T_REG_ is shown. (H) Relative mRNA expression of *TP53BP2, TNFSF11* and *IL23* in human primary CD4^+^ T_REG_ separated by FACS from PBMCs (n = 4, patients 157, 801, 673, 366, Supplementary Table S1) and plated in 6-well plates alone or with 10 μg/mL of CD19-4-1BBL for 22 h. Three technical replicates were analyzed for each sample. Data were normalized to β-actin expression. Gene expression was determined by calculating the difference (ΔCt) between the threshold cycle (Ct) of each gene and that of the reference gene and was expressed as the mean of 3 replicates ± SEM. Then the relative quantification values were calculated as the fold change expression of the gene of interest over its expression in the selected cell type reference sample, i.e., the untreated sample (considered as the calibrator sample), by the formula 2 ^-ΔΔCt^. Finally, the treated sample relative mRNA expression was normalized to the vehicle. Samples with undetermined Ct values were not included.

### CD19-4-1BBL TARGETS IMMUNOSUPPRESSIVE CD4^+^ FOXP3^+^ T_REG_ CELLS

Unlike memory T cells, the role of 4-1BB-4-1BBL costimulation on T_REG_ cells is unclear and, to some extent, controversial^22,23^. We first demonstrated in cell depletion studies that, in half of the CLL patient samples analyzed, CD19-4-1BBL bispecific molecule depleted CD4^+^ FoxP3 T_REG_ cells (**Figure 2D**). When we analyzed the transcriptome of T_REG_ cells exposed to CD19-4-1BBL in vitro (**Figure 1A**), we observed the coexistence of signals of cell activation and death. We found several pathways related to the 4-1BB signalosome, including the p38/MAP kinase signaling pathway (**Figure 2E**) and GO analysis enriched terms related to the 4-1BB signalosome, including NF-κB binding and BCL2-mediated lymphocyte proliferation (**Figure 2F**). Of note, in both pathway and GO analyses, we also found upregulated the apoptotic signaling and functions related to the suppression of T_REG_ cell activity, including anoikis and apoptotic cell clearance (**Figures 2E, F**). Like memory T cells, we identified several upregulated pathways related to infection immunity, including the regulation of viral transcription and of viral entry into host cells (**Figure 2F**). Our data agree with published studies demonstrating that 4-1BB agonistic therapy can reprogram T_REGS_ into cytotoxic T cells with anti-tumor activity.^23^ A supervised heatmap for selected, differentially expressed genes is described in **Figure 2G**. We validated by RT-PCR crucial mediators of T_REG_ suppressive function, including TP53BP2, IL23A and TNFSF11 (**Figures 2G, H**). TP53BP2, also known as Bcl-2 binding protein, plays a central role in the regulation of apoptosis and cell growth. It was found upregulated together with BCL2 transcript. Finally, we evaluated the expression level of selected proteins via full spectrum flow cytometry on an independent cohort of CLL patient samples (Table S1, **Figure S3A**). Upon CD19-4-1BBL, we confirmed the depletion of CD4^+^ T_REG_ cells and we found that TNFSF11 was significantly downregulated, while IL23A was upregulated in half of the samples analyzed (**Figure S3B-E).**

### CD19-4-1BBL SHARPENS THE TRANSCRIPTOME OF MONOCYTES AND MDSCs

Upon treatment with the 4-1BB-4-1BBL bispecific monocyte subsets and M-MDSCs underwent drastic changes in their transcriptomes. As for I monocytes, the pathway analysis highlighted the upregulation of the interferon and T-cell related activation signaling (**Figures 1A, 3A**). The GO analysis on I monocytes reported enriched terms related to cytokine receptor binding, interferon gamma-mediated signalling, regulation of lymphocyte activation, and positive regulation of neutrophil function (**Figure 3B**). A number of enriched terms and pathways were involved in the immune response to infections. Overall, the transcriptome of I monocytes was profoundly modified through the modulation of several genes involved in monocyte function, monocyte/macrophage differentiation and response to infections. **Figure 3C** shows a supervised analysis of differentially expressed transcripts in this monocyte subset. Several of these transcripts were validated by RT-PCR, including SELL/CD62L, which encodes for the selectin CD62L normally expressed on circulating C monocytes with antimicrobial function and able to differentiate into antitumor macrophages^24^ (**Figure 3D**).

**Fig. 3.**
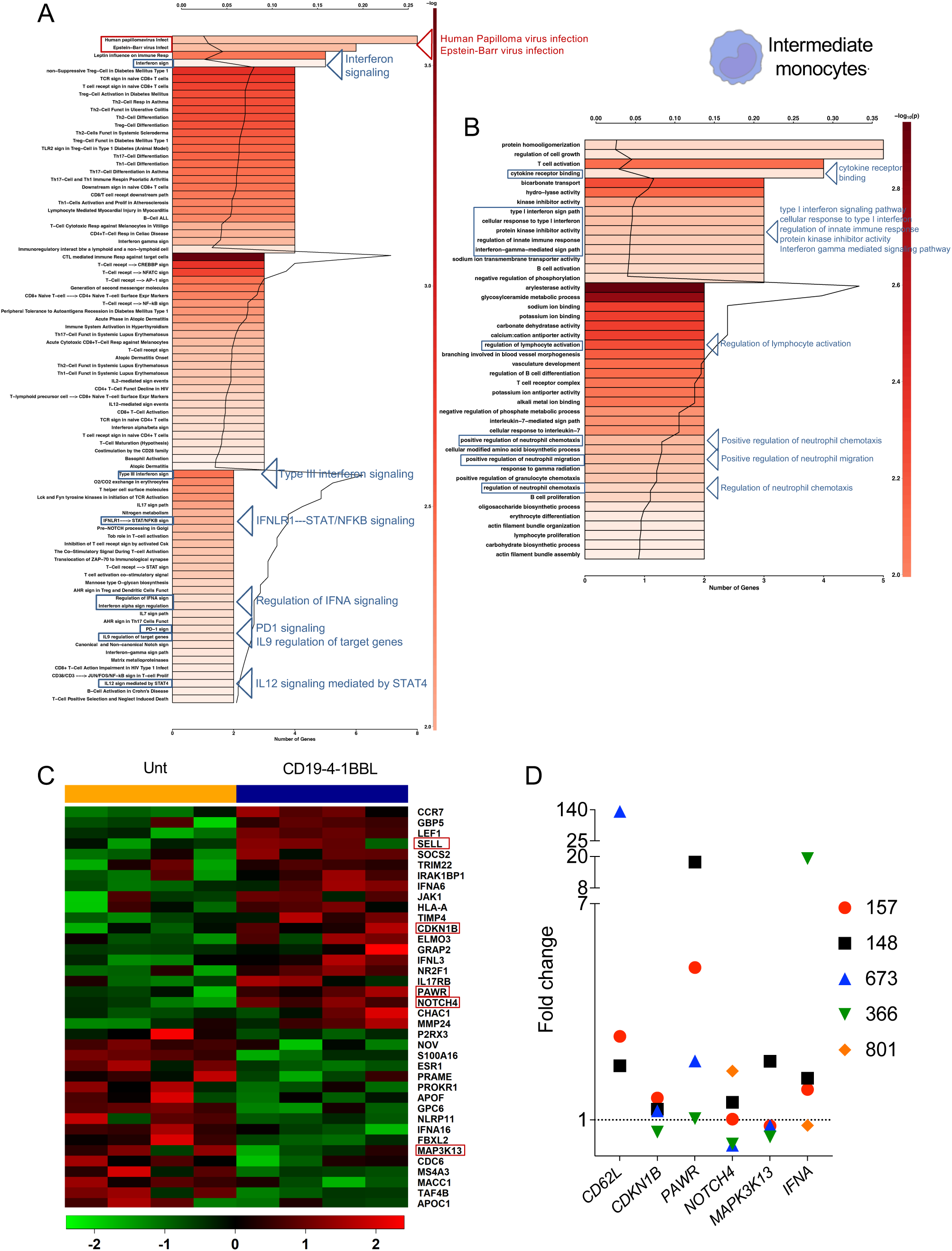
Transcriptome analysis of patient-derived intermediate monocytes. (A) GO and (B) pathway analyses of differentially expressed transcripts between CD19-41BBL treated and untreated I monocytes are shown. (C) A supervised heatmap for chosen, differentially expressed transcripts on Intermediate monocytes is shown. (D) Relative mRNA expression of *CD62L, CDKN1B, PAWR, NOTCH4, MAPK3K13* and *IFN*α in human primary I monocytes separated by FACS from PBMCs (n = 5, patients 157, 148, 673, 366, 801, Supplementary Table S1) and plated in 6-well plates alone or with 10 μg/mL of CD19-4-1BBL for 22 h. Three technical replicates were analyzed for each sample. Data were normalized to β-actin expression. Gene expression was determined by calculating the difference (ΔCt) between the threshold cycle (Ct) of each gene and that of the reference gene and was expressed as the mean of 3 replicates ± SEM. Then the relative quantification values were calculated as the fold change expression of the gene of interest over its expression in the selected cell type reference sample, i.e., the untreated sample (considered as the calibrator sample), by the formula 2 ^-ΔΔCt^. Finally, the treated sample relative mRNA expression was normalized to the vehicle. Samples with undetermined Ct values were not included.

When we analyzed C monocytes, we found that PI3K/AKT and FGFR1-2 signaling were activated (**Figures 1A, 4A**). GO analysis showed the activation of the 4-1BB signalosome-related regulation of MAPK activity and the enrichment of a number of terms, including the positive regulation of intracellular signal transduction, cellular response to cytokine stimulus, positive regulation of T-helper 1 immune response and of IL2 biosynthesis process (**Figure 4B**). Of note, we found that terms such as the defence response to fungus were enriched in this monocyte subset. We selected a number of differentially expressed transcripts depicted in the heatmap (**Figure 4C**). Many transcripts involved in monocyte function, differentiation into macrophages or dendritic cells (GAS6, CD24), and macrophage function polarization (MCOLN2, EIF4E), were validated by RT-PCR (**Figure 4D**). CD24 is expressed on hematopoietic cells, including dendritic cells (DC) and macrophages. Distinct DC subsets dictate the fate decision between effector and central memory CD8^+^ T cell differentiation by a CD24-depepndent mechanism^25^. IRF4 mRNA is normally upregulated during DC differentiation^26^ and was found upregulated on C monocytes upon CD19-4-1BBL treatment (**Figure 4D**).

**Fig. 4.**
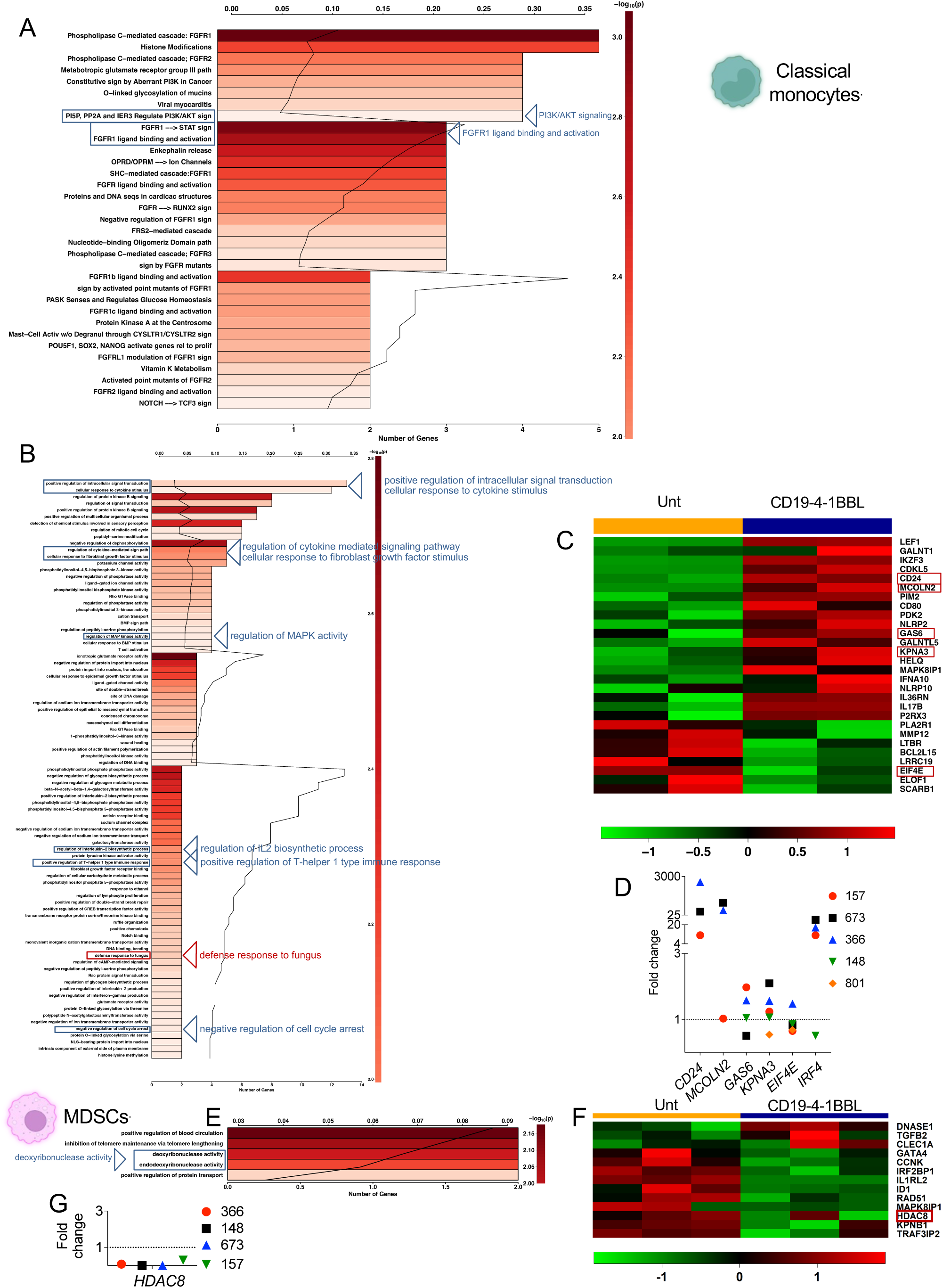
Transcriptome analysis of patient-derived classical monocytes and monocytic myeloid-derived suppressor cells. (A) GO and (B) pathway analyses of differentially expressed transcripts between CD19-41BBL treated and untreated I monocytes are shown. (C) A supervised heatmap for chosen differentially expressed transcripts on Classical monocytes is shown. (D) Relative mRNA expression of *CD24, MCOLN2, GAS6, KPNA3, EIF4E* and *IRF4* in human primary C monocytes separated by FACS from PBMCs (n = 5, patients 157, 148, 673, 366, 801, Supplementary Table S1) and plated in 6-well plates alone or with 10 μg/ml of CD19-4-1BBL for 22h. Three technical replicates were analyzed for each sample. Data were normalized to β-actin expression. Gene expression was determined by calculating the difference (ΔCt) between the threshold cycle (Ct) of each gene and that of the reference gene and was expressed as the mean of 3 replicates ± SEM. Then the relative quantification values were calculated as the fold change expression of the gene of interest over its expression in the selected cell type reference sample, i.e., the untreated sample (considered as the calibrator sample), by the formula 2 ^-ΔΔCt^. Finally the treated sample relative mRNA expression was normalized to the vehicle. Samples with undetermined Ct values were not included. (E) GO analysis of differentially expressed transcripts between CD19-41BBL treated and untreated M-MDSCs is shown. (F) A supervised heatmap for chosen differentially expressed transcripts on M-MDSCs is shown. (G) Relative mRNA expression of *HDAC8* in human primary M-MDSCs separated by FACS from PBMCs (n=4, patients 157, 148, 673, 366, Supplementary Table S1) and plated in 6-well plates alone or with 10 μg/mL of CD19-4-1BBL for 22 h. Three technical replicates were analyzed for each sample. Data were normalized to β-actin expression. Gene expression was determined by calculating the difference (ΔCt) between the threshold cycle (Ct) of each gene and that of the reference gene and was expressed as the mean of 3 replicates ± SEM. Then the relative quantification values were calculated as the fold change expression of the gene of interest over its expression in the selected cell type reference sample, i.e., the untreated sample (considered as the calibrator sample), by the formula 2 ^-ΔΔCt^. Finally, the treated sample relative mRNA expression was normalized to the vehicle. Samples with undetermined Ct values were not included.

Finally, we analyzed the transcriptome of M-MDSCs. Unlike monocytes, the pathway analysis of MDSCs showed a high number of downmodulated pathways, including the nitric oxide stimulation pathway, the TGF-beta receptor complex signalling pathway and the mechanism of protein import into the nucleus (**Figure S4A**) and, of note, the upmodulation of pathways involved in T cell activation, co-stimulation and NF-κB signaling (**Figure S4B**). The GO analysis showed the enrichment of terms such as the deoxyribonuclease and endodeoxyribonuclease activities that are involved in the DNA fragmentation during apoptosis (**Figure 1A, 4E**), while the majority of cellular and molecular functions were found downmodulated, including those for protein-kinase binding, cellular response to DNA damage stimulus, regulation of signal transduction, and cell-cycle process, DNA replication, DNA polymerase activity, protein localization to cell surface (**Figure S5**). Relevant transcripts were selected and described in the heatmap (**Figure 4F**). HDAC8 was found downmodulated on M-MDSCs upon CD19-4-1BBL treatment and was validated by RT-PCR in all samples analyzed (**Figure 4G**). HDAC8 encodes for a protein belonging to the class I histone deacetylase family, it has histone-deacetylase activity and represses transcription when tethered to a promoter. ^27^ Inhibition of class I and class II HDAC induce inactivation of MDSCs.^28^ Taken together, these findings indicate that the 4-1BB signalosome profoundly shapes the transcriptome of monocytes and M-MDSCs.

### CD19-4-1BBL CONFERS AN ANTITUMOR IMMUNOPHENOTYPE TO MYELOID CELLS

When the PBMCs of patients with CLL were treated in vitro with CD19-4-1BBL, we observed a dose-dependent, antileukemic effect. ROR1 is associated with progressive CLL,^29^ and we used it to identify CLL cells and to avoid flow cytometry CD19-related competitive binding reactions (**Figure 5A**). Next, we wondered whether myeloid cells in the non-malignant microenvironment with known protumor function (e.g. M-MDSCs and tumor-associated macrophages) might be involved in the antileukemic effect observed in vitro upon CD19-4-1BBL treatment. We designed and optimized a 32-marker panel for high-dimensional, full spectrum flow cytometry, including surface markers, transcription factors, cytokines, chemokines, and apoptotic proteins, among others (**Figure 5B**). We validated the upregulation of CD62L selectin on intermediate monocytes in all the patients analyzed (**Figures 5C-D**). Since this selectin is normally expressed on classical monocytes with anti-microbial function that differentiate into antitumor macrophages, these findings confirm our hypothesis that 4-1BB-signalosome stimulation on monocytes has a role in monocyte differentiation and triggers their antitumor function. Though not reaching statistical significance, interferon (IFN) was found upregulated on intermediate monocytes (**Figures 5E-F**) in 6/10 samples, thus confirming the transcriptional activation of the IFN signaling pathway (**Figures 3A-D**). Previous evidence demonstrated that selected DC subsets dictate the fate decision of effector and central memory T cell differentiation through a CD24-dependent mechanism. Of interest, we have found CD24 upregulated on classical monocytes in half of the patients analyzed (**Figures 5G-H**). Since this molecule is normally expressed on DCs, our data demonstrate that CD19-4-1BBL stimulates the differentiation of classical monocytes into monocytic-derived DCs. In addition, EIF4E was found downmodulated on classical monocytes (**Figures 5I-J**) in most samples.

**Fig. 5.**
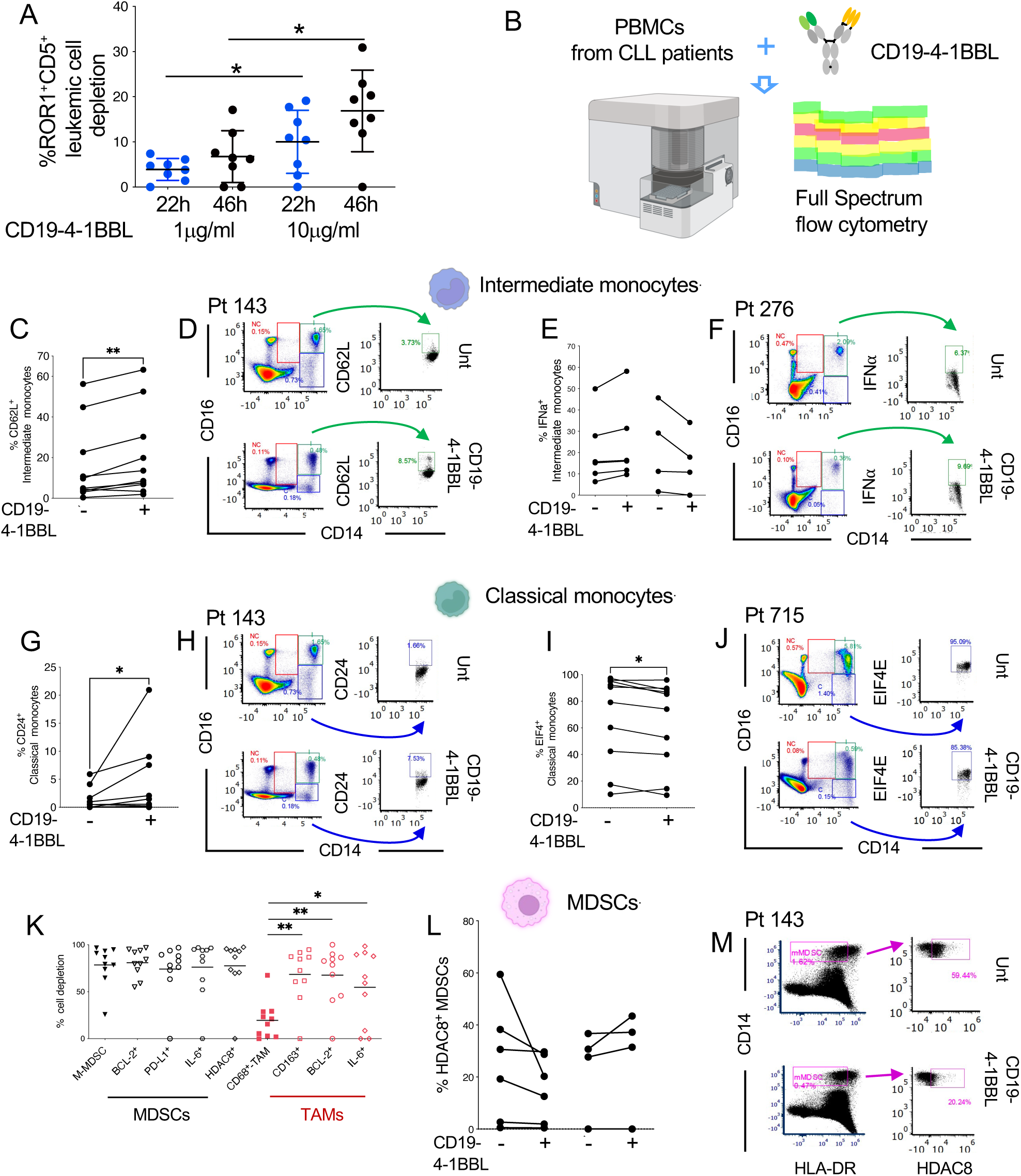
Immunophenotype of myeloid cells upon CD19-4-1BBL treatment. (A) Leukemic CD19^+^ ROR1^+^ CD5^+^ cell depletion (n = 25, Supplementary Table S2) after 22 h and 46 h of treatment with 1µg/mL and 10µg/mL of CD19-4-1BBL was analyzed by multi-color flow cytometry and calculated by the formula: the following formula: 100 - % remaining cells, where % remaining cells = (absolute number in treated samples/absolute number in untreated samples) × 100. A statistical analysis was performed using the Student *t* test \**P* < 0.05. Details are described in the Supplementary Methods. (B) Fresh PBMCs from CLL patients (n = 10) were incubated with CD19-4-1BBL (10µg/mL, Roche) for 48 h and then stained and analyzed by spectral flow cytometry. (C) the relative contribution of CD62L^+^ CD14^++^ CD16^+^ intermediate monocytes treated with CD19-4-1BBL or left untreated and (D) related dot plots from representative patient 143 are shown. *P* value is given by Mann-Whitney-Wilcoxon test \*\**P* < 0.01. (E) The relative contribution of IFNα^+^ CD14^++^ CD16^+^ intermediate monocytes from patients with CLL (n=10, Table S1) treated with CD19-4-1BBL or left untreated and (F) related dot plots from representative patient 276 (Table S1) are shown. (G) The relative contribution of CD24^+^ CD14^++^ CD16^-^ classical monocytes from patients with CLL (n = 10, Table S1) treated with CD19-4-1BBL or left untreated and (H) related dot plots from representative patient 143 are shown. *P* value is given by Mann-Whitney-Wilcoxon test \**P* < 0.05. (I) The relative contribution of EIF4^+^ CD14^++^ CD16^-^ classical monocytes from patients with CLL (n=10, Table S1) treated with CD19-4-1BBL or left untreated and (J) related dot plots from representative patient 715 are shown. *P* value is given by Mann-Whitney-Wilcoxon test \**P* < 0.05. (K) CD14^+^HLADR^low/-^ M-MDSC and Lin^-^ CD68^+^ TAM cell depletion (n = 10, Table S1) after 48h of treatment with 10µg/ml of CD19-4-1BBL was analyzed by spectral flow cytometry and calculated by the formula: 100 - % remaining cells, where % remaining cells = (absolute number in treated samples/absolute number in untreated samples) ×100. *P* value is given by Mann-Whitney-Wilcoxon test \**P* < 0.05, \*\**P* < 0.01. Details are described in the Supplementary Methods. (L) The relative contribution of CD14^+^HLADR^low/-^ HDAC8^+^ M-MDSC from patients with CLL (n = 10, Table S1) treated with CD19-4-1BBL or left untreated and (M) related dot plots from representative patient 143 are shown. *P* value is given by Mann-Whitney-Wilcoxon test \**P* < 0.05, \*\**P* < 0.01.

Upon CD19-4-1BBL treatment, we observed the depletion of M-MDSCs expressing BCL2, PD-L1, IL6 and HDAC8 (**Figure 5K**). HDAC8 was found downmodulated on MDSCs in 6 of 10 samples (**Figures 5L-M**), but it did not reach statistical significance due to the heterogeneity of the data. We recently demonstrated that monocytes and MDSCs from patients with CLL differentiate into TAMs.^7^ Of note, when we analyzed human CD68^+^ macrophages spontaneously differentiating in vitro in PBMCs cultures (**Figures 6A**), we confirmed depletion of CD68^+^ CD163^+^ and CD68^+^ CD206^+^ protumor-associated macrophages upon CD19-4-1BBL treatment (**Figures 5K, 6A-C**). Of note, we found CD68^+^ CD80^+^ and CD68^+^ CD86^+^ antitumor-associated macrophages in PBMCs cultures upon CD19-4-1BBL treatment (**Figures 6D)**.

**Fig. 6.**
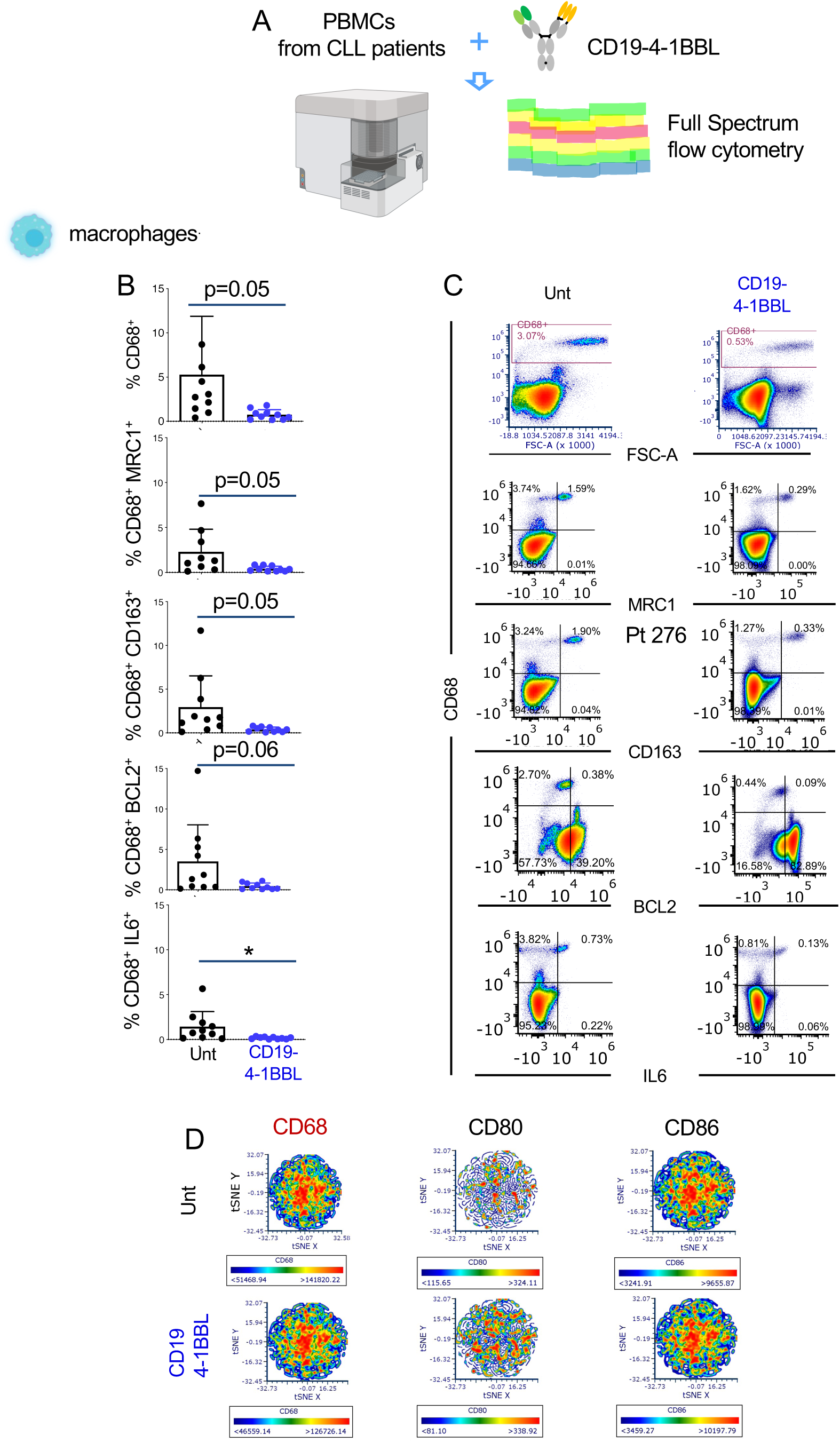
Patient-derived macrophage characterization upon CD19-4-1BBL treatment. Fresh PBMCs from CLL patients (n = 10, table S1) were incubated with CD19-4-1BBL (10 g/mL, Roche) for 48 h and then stained and analyzed by spectral flow cytometry. (B) The mean values ± SDs of the relative contributions of the whole pool of CD68^+^ TAMs and MRC1, CD163, BCL2, IL6 expressing CD68^+^ TAMs and (C) related representative plots are shown. (D) tSNE plots (pre-gated on CD68^+^ cells) representing CD68^+^, CD80^+^ and CD86^+^ fluorescence intensity of untreated and CD19-4-1BBL-treated cells from 10 patient samples are shown. A statistical analysis was performed using the Student *t* test \**P* < 0.05.

### CD19-4-1BBL AND CD20 OBINUTUZUMAB ERADICATE CLL IN A HUMANIZED, PATIENT-DERIVED XENOGRAFT MODEL

We successfully established an innovative patient-derived xenograft (PDX) system of CLL based on the humanized MISTRG mouse model. This mouse strain was developed to facilitate the engraftment of the entire human immune system, including myeloid cells^17^. Human myeloid cells (NC, INT and C monocytes and M-MDSCs) and lymphoid cells (T_EM_ and T_CM_ CD8^+^ T cells and CD4^+^ T_REG_) from the PB of CLL patients were adoptively transferred into MISTRG mice preinjected with MEC1 cells. MEC1 is a cell line derived from a patient with CLL in prolymphocytic transformation (PLL), and mice transplanted with this cell line recapitulate aggressive CLL/PLL.^30^

The MISTRG-based PDX system was used to evaluate the immune-based antileukemic activity of the bispecific molecule CD19-4-1BBL in vivo. After immune reconstitution, MEC1-xenotransplanted mice, were treated with the bispecific CD19-4-1BBL as monotherapy or in combination with CD20 obinutuzumab (**Figure 7A**). PDX immune reconstitution was achieved by transplanting mice with human immune cells separated from the PBMCs of patients with either stable (**Figures 7B-C**) or high risk/progressive CLL (**Figures 7D-E**). The novel combination favorably improved the survival in 3 separate experiments (**Figures 7B-G**) and cured some of the mice (**Figure S6**). A better survival outcome was observed in mice reconstituted with immune cells from patients with high-risk CLL (**Figures 7D-G, Table S22**). Of note, in one of the experiments, treatment started at day 21, when mice showed severe signs of leukemia (**Figures 7F-G**).

**Fig. 7.**
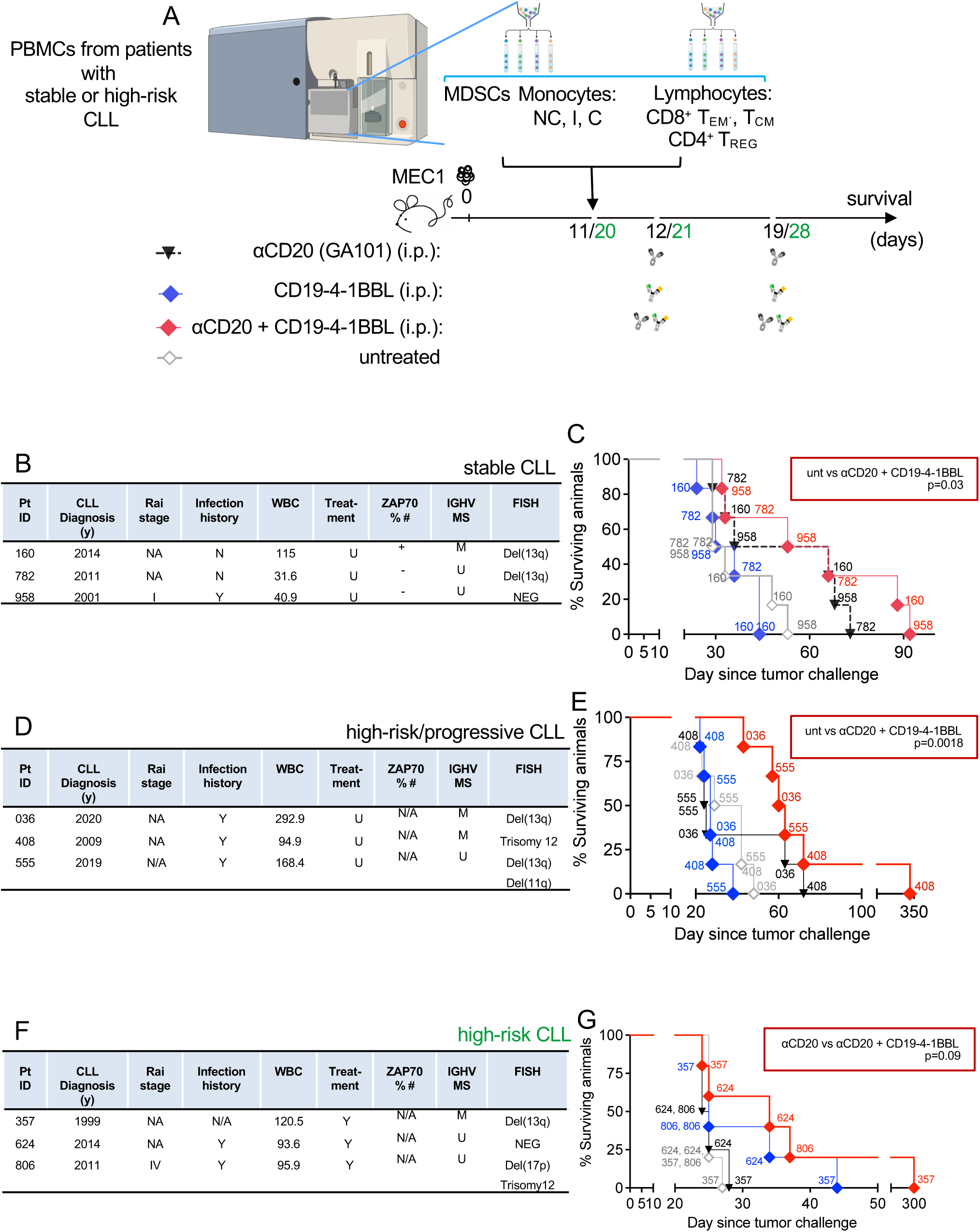
Anti-leukemic effect and survival impact of CD19-4-1BBL treatment in the MISTRG-based xenograft system. (A) MISTRG mice transplanted intravenously (i.v.) with MEC1 cells (day 0) received weekly injections of CD19-4-1BBL (1mg/kg, blue rombi) or CD20 (10mg/kg, black triangles) or their combination (red rombi), after the adoptive transfer of patient-derived immune cells (days 11 or day 20, depending on the experiment). The transferred cells include NC (682-6203 cells depending on the patient), I (305-43354 cells depending on the patient), C (4234-256552 cells depending on the patient) monocyte subsets, M-MDSCs (174-63051 cells depending on the patient), CD4^+^ T_REG_ (4521-27174 cell depending on the patient), CD8^+^ effector T_EM_ (6457-61364 cells depending on the patient) and central memory T_CM_ (2738-25807 cells depending on the patient) cells separated from the PBMCs of patients described in Tables B, D, F using fluorescence-activated cell sorting. Mice were monitored for survival in 3 independent experiments (n=6 mice/group). (B-C) Clinical features of related patients with stable CLL and Kaplan-Meyer survival curve are shown. (D-E) Clinical features of related patients with high-risk/progressive CLL and Kaplan-Meyer survival curves are represented. (F-G) Clinical features of related patients with high-risk CLL and Kaplan-Meyer survival curves are represented.

To evaluate the effect of CD19-4-1BBL bispecific as monotherapy or in combination with CD20 on patient-derived immune cells, we sacrificed immune-reconstituted MEC1-xenotransplanted mice 6-7 days after the last treatment in a separate experiment (**Figures S7A-B**). The combination favorably reduced circulating CD23^+^ leukemic cells (**Figure S7C**) and selectively targeted protumor immune cells from patients with high-risk CLL in the peripheral blood (PB) and bone marrow (BM), including CD8^+^ PD1^+^ T_CM,_ M-MDSCs and CD163^+^ TAMs (**Figures S8B-G**). Of note, we observed a significant reduction of circulating CD4^+^ FoxP3 T_REG_ cells upon CD19-4-1BBL (**Figure S8D**), that was not accompanied by a significant increase of memory or naïve CD4^+^ T cells (**Figure S8E**). This evidence confirms our transcriptomic and in vitro cell depletion data on CD19-4-1BBL-induced targeting of T_REG_ cells (**Figures 2-S3**). To final support the conclusion that CD19-4-1BBL mitigates the suppressive activity of the remaining T_REG_ cells and M-MDSCs, we performed in vitro suppression assays with patient-derived immune cells. In two out of three samples analyzed, we observed a reduced suppressive activity of CD4^+^ FoxP3^+^ T_REG_ and MDSCs on CD8^+^ naïve and T_CM_ lymphocytes, in terms of IFN production and proliferation (**Figures S9-S10**).

Altogether, these data suggest that CD19-4-1BBL in combination with CD20 obinutuzumab confers antitumor function to patient-derived immune cells and improves the survival. Of interest, the highest antitumor efficacy and leukemia eradication was mediated by immune cells originating from high-risk/progressive CLL, thus highlighting the rationale to use CD19-4-1BBL bispecific immunomodulator in patients with high-risk/progressive disease.

## DISCUSSION

Tumor-targeting costimulatory bispecific antibodies have gained great interest in immunotherapy given their ability to exert multiple mechanisms of action at the same time.^31^ Nine bispecific antibodies have been approved for solid tumors and hematologic malignancies, including acute lymphoblastic leukemia, multiple myeloma, and diffuse large B cell lymphoma. However, no bispecific antibodies are currently clinically approved for the treatment of CLL.

In this study, we investigated for the first time the antileukemic activity of the CD19-4-1BBL bispecific molecule in CLL, and we identified multiple mechanisms of action involving cells of the adaptive and innate immunity. Multiomics studies demonstrated that, upon CD19-4-1BBL treatment, selected non-malignant cells of either the myeloid and lymphoid lineage rapidly undergo transcriptomic and phenotypic modifications. The antitumor immunophenotype found in vitro aligned with the antileukemic function we observed in survival studies performed in MISTRG-based PDX mice.

4-1BB is a costimulatory receptor very well studied on T lymphocytes and NK cells, and its function has been extensively exploited in cancer immunotherapy. It is also expressed on myeloid cells, including monocytes and macrophages, where its activity still remains elusive.^32^ Several 4-1BB agonist antibodies, including urelumab^33^ and utomilumab,^34,35^ have been tested in early-phase clinical trials. Severe liver toxicity has been observed in patients treated with urelumab likely due to 4-1BB cross-linking via FcγRIIB.^36^ A lack of efficacy has been reported in patients treated with utomilumab.^33^ These clinical observations helped the design of next generation 4-1BB agonists with higher efficacy and reduced liver toxicity.^37^ 4-1BB agonists have been implemented by combining an agonistic 4-1BB binding site with a second target.^37^ The second target includes a surface tumor-associated antigen such as HER2^38^, EGFR^39^, or fibroblast activating protein alpha (FAP).^40^ Of note, the CD19-41BBL bispecific fusion protein has been recently developed with modifications of the Fc region that abrogate FcγR-mediated cross-linking. The combination of CD19-41BBL with T-cell bispecific antibodies has been reported to eradicate tumors in mice bearing aggressive human lymphoma^8^. Claus *et al* demonstrated that crosslinked CD19-4-1BBL induces in vitro T-cell activation,^8^ but its activity on myeloid cells has not been explored. 4-1BB relies on TRAFs to create the 4-1BB-4-1BBL signalosome.^13^ Our transcriptome studies on PBMCs from patients with CLL confirmed that the 4-1BB-4-1BBL costimulation of T cells activates the downstream NF-kB/JNK pathways through selected TRAFs.^41,13,14^ Additionally, our transcriptome and TCR-sequencing studies on memory CD8^+^ T cells demonstrated the induction of a T-cell response machinery to viral infections and virus-specific T-cell clonotypes, thus confirming the role of 4-1BB-4-1BBL costimulation in the antiviral immunity.^14^

The effect of 4-1BB stimulation on CD4^+^ T_REG_ cells is controversial. 4-1BB is expressed on T_REG_ cells,^42^ and properly engineered anti-4-1BB monoclonal antibodies have been preclinically tested to selectively deplete these immunosuppressive cells.^22^ Of interest, 4-1BB costimulation on T_REGS_ augments their proliferation without enhancing their immunosuppressive function.^43,44^ Therapeutic 4-1BB costimulatory signaling can reprogram T_REGS_ into cytotoxic effector T cells with antitumor function.^23^ In line with this evidence, upon CD19-4-1BBL treatment, we observed either the depletion or the activation of the 4-1BB-4-1BBL signalosome machinery and stimulation of the T-cell proliferation pathways on CD4^+^ T_REG_ cells from patients with CLL. CD4^+^ T_REG_ cell depletion might involve other immune cells, and this should be addressed in future studies. In line with previous studies on colorectal cancer patients^45^, we found TNFS11, also known as RANKL, upregulated on CD4^+^ T_REG_ cells from patients with CLL. We demonstrated that the CD19-4-1BBL bispecific molecule targets the immunosuppressive activity of CD4^+^ T_REG_ cells through the RANKL signaling pathway, upon the 4-1BB-4-1BBL signalosome activation. Overall these data support the dual effect of 4-1BB costimulation on T_REG_ cells, which was previously described in several studies.^22,42,43,23^

It has been shown that the 4-1BB-4-1BBL costimulation activates monocytes, and increases their proliferation and survival as well as the release of proinflammatory cytokines such as TNFα and IL8.^46,47^ The crosslinking of 4-1BB on monocytes induces the apoptosis of B lymphocytes, thus playing a role in the monocyte-dependent control of B-cell survival.^48^

We demonstrated that CLL patient-derived monocytes and M-MDSCs treated with 4-1BB-4-1BBL bispecific undergo deep changes of their transcripts involved in monocyte function, differentiation into macrophages, macrophage polarization and response to infections. Our studies align with the evidence on 4-1BB-related metabolic and functional reprogramming of monocytes and macrophages^49^ and on the induction of the differentiation of CD34^+^ cells into monocytes upon 4-1BB ligand signaling.^50^ We investigated for the first time the agonistic activity of the CD19-4-1BBL fusion molecule on M-MDSCs, and we observed the depletion of these protumor myeloid cells. Of note, HDAC8 was found downmodulated on M-MDSCs due to CD19-4-1BBL treatment. HDAC8 is a histone deacetylase enzyme involved in cancer cell proliferation, immune evasion, and drug resistance^51^ and is considered as a promising therapeutic target.^52^ Our data highlight the crucial contribution of HDAC8 to the M-MDSCs-related mechanism of action of the CD19-4-1BBL fusion molecule. We recently demonstrated that patient-derived monocytes and M-MDSCs differentiate into protumor TAMs.^7^ Protumor TAMs were significantly depleted in our in vitro and in vivo studies, thus confirming the previously observed role of 4-1BB-4-1BBL costimulation in monocyte/macrophage differentiation.

Intriguingly, our in vivo xenograft studies highlighted the crucial antileukemic contribution of immune cells originating from patients with high-risk CLL upon the treatment with the combination of CD19-4-1BBL and αCD20 obinutuzumab. These findings prompt us to speculate that the 4-1BB-crosslinking activity on adaptive and innate immune cells from patients with CLL might change depending on the disease features. We hope that our studies will pave the way for future clinical trials of CD19-4-1BBL in patients with CLL.

## Author contributions

P.K. performed experiments, analyzed the data and helped writing the manuscript; R.Z. performed experiments and analyzed the data; C.I. performed the statistical analysis of the data; S.S. assisted in spectral flow cytometry studies; P.B. performed cell depletion experiments; N.S. performed the TCR sequencing data analysis; H.P. performed immunohistochemistry; V.J. assisted in mice studies; K.C-K. assisted in flow cytometry and cell sorting study design, set up and analysis in mice and human experiments; W.W. contributed patient samples with clinical/biological features and assisted in the interpretation and discussion of the results; M.T.S.B. designed and supervised the project, performed experiments, analyzed the data and wrote the paper. All the co-authors reviewed the manuscript.

## Supporting information

Khare P et al Supplemental Materials

Khare P et al Table S7

Khare P et al Table S8

Khare P et al Table S9

Khare P et al Table S10

Khare P et al Table S11

Khare P et al Table S14

Khare P et al Table S15

Khare P et al Table S6

Khare P et al Table S17

Khare P et al Table S18

Khare P et al Table S19

Khare P et al Table S20

Khare P et al Table S21

## Acknowledgments

We are particularly grateful to Michael J Keating for his extraordinary contribution to CLL research. We are grateful to the DVMS Veterinary Pathology Services at UT MD Anderson for the immunohistochemical studies. We are grateful to Christian Klein and Sylvia Herter for providing constructive comments and crucial suggestions on CD19-4-1BBL, particularly on the in vitro experiments. Some of the figures were created using BioRender.com. and Servier Medical Art: www.smart.servier.com. We are grateful to Laura Russell and the Department of Scientific Publications, MD Anderson Cancer Center, for reviewing the manuscript and providing constructive comments.

