## Supplementary material for "The CD19-4-1BBL antibody fusion protein unleashes the immune system against high-risk chronic lymphocytic leukemia": Khare P et al Supplemental Materials

**The PDF file includes:**

Supplemental Materials and Methods
References for Supplemental Methods
Tables S1 to S6
Legends for Tables S7 to S11
Tables S12-S13
Legends for Tables S14 to S21
Table S22
Figs. and Legends S1 to S10
References for Table S11

#### SUPPLEMENTARY MATERIALS AND METHODS

##### Cell lines

MEC1 cell line is a CD5$^{low/-}$ CLL cell line established from a CLL patient in prolymphocytoid transformation to B-cell prolymphocytic leukemia (B-PLL). It was cultured in RPMI 1640 medium (Invitrogen, Carlsbad, CA, USA) with 10% fetal bovine serum and gentamicin (15 µg/mL; Sigma-Aldrich, St. Louis, MO, USA). MEC1 cell line regularly tested negative for *Mycoplasma* contamination.

##### Conventional flow cytometry-based cell-depletion assay

For cell depletion studies, fresh PBMCs from untreated CLL patients were seeded, as triplicates, at 3 x $10^6$ cells/mL in culture medium and treated with CD19-4-1BBL (1 µg/mL, 10 µg/ml) or RPMI vehicle for 22 h and 46 h. The specific percentages of remaining cells in the treated samples were calculated as (the absolute number of cells in treated samples/the absolute number of cells in control samples) x 100. For each condition, the absolute number of remaining cells was calculated as the total number of viable cells (trypan blue exclusion determination) x the percentage of viable cells (flow cytometry analysis determination). Then, specific cell depletion was calculated as follows: 100 - the specific percentage of remaining cells, as described [1]. The flow cytometry analysis (Fortessa X-20, BD Biosciences) of human myeloid and lymphoid cell types is described below.

8-color flow cytometry phenotype analysis of human live myeloid cells and 11-color flow cytometry phenotype analysis of human live lymphoid cells were performed using

LSRFortessa X-20 with the antibodies described in Supplementary Tables S3-S4. PBMCs were first incubated with LIVE/DEAD fixable Aqua dye; then, after the blocking of Fc receptors, the cells were stained with the surface antibodies described in Supplementary Tables S3-S4. Finally, the cells were incubated with ammonium chloride solution (STEMCELL Technologies) to lyse red cells. For lymphoid cell Foxp3 detection, surface-stained cells were further fixed and permeabilized using a $T_{REG}$ detection Kit (Miltenyi Biotec, Bergisch Gladbach, Germany) and finally stained with an anti-Foxp3 antibody.

**Human cell sorting**

For the TCR-sequencing, transcriptional, and patient-derived-xenograft studies human myeloid and lymphoid cells were separated by fluorescence-activated cell sorting.

Live myeloid cells were isolated by 4-way, fluorescence-activated cell sorting, after surface staining with the following antibodies: Alexa Fluor 700 mouse anti-human CD66b (G10F5), APC mouse anti-human Lineage Cocktail (CD3/CD19/CD20/CD56) (UCHTI, HIB19, 2H7, 5.1H11), Brilliant Violet 786 mouse anti-human CD14 (MφP9) purchased by BD Biosciences, PE mouse anti-human CD16 (3G8) purchased by Biolegend (San Diego, CA, USA), APC-Cy7 mouse anti-human HLA DR (L243).

Live lymphoid cells were isolated by 4-way fluorescence-activated cell sorting after surface staining with the following antibodies: eFluor 450 mouse anti-human CD8a (SK1) purchased by eBiosciences (Waltham, MA, USA), PerCP mouse anti-human CD45RO (UCHL1) purchased by Biolegend, PE-Cy7 mouse anti-human CD45RA (L48) purchased by BD Biosciences, Brilliant Violet 605 mouse anti-human CD62L (DREG-56), PE-Dazzle 594 mouse anti-human CD4 (SK3), PE mouse anti-human CD25 (4E3) purchased by Milteny Biotec, APC mouse anti-human CD19 (J3-119) purchased by Beckman Coulter, FITC mouse anti-human CD127 (HIL-7R-M21) purchased by BD Biosciences.

Live/Dead Fixable Aqua (Thermo Fisher Scientific) staining was first performed to allow the discrimination of Live/Dead cells.

The identification of the human immune cell populations was performed as we previously published[2]. Monocyte subsets were identified using a negative exclusion gating strategy. We excluded CD66b$^{+}$ neutrophils, then, using a lineage (Lin) cocktail including mAbs to CD3, CD19, CD20, and CD56, we excluded T cells, B cells, and NK cells, respectively. CD14 and CD16 expression was then used to identify

$CD14^{+}CD16^{++}$ nonclassical (NC), $CD14^{++}CD16^{+}$ intermediate (I), and $CD14^{++}CD16^{-}$ classical (C) monocytes. M-MDSCs were identified as $CD14^{+}HLA\text{-}DR^{low/-}$.

T cells were identified as follows: naïve $hCD8^{+}CD45RO^{-}CD45RA^{+}CD62L^{+}$ T cells; central memory $hCD8^{+}CD45RA^{-}CD45RO^{+}CD62L^{+}$ $T_{CM}$, effector memory $hCD8^{+}CD45RA^{-}CD45RO^{+}CD62L^{-}$ $T_{EM}$ and $hCD4^{+}CD25^{+}CD127^{low}$ $T_{REG}$.

**T-cell suppression assays**

The suppressive capacity of patient-derived $T_{REGS}$ and M-MDSCs was tested as previously described[3, 4]. PBMCs from CLL patients were seeded in culture medium and treated with CD19-4-1BBL (10 µg/ml) or RPMI vehicle for 24 h. In the $T_{REG}$ suppression assay, fluorescent-activated cell sorted $CD4^{+}CD25^{+}CD127^{low/-}$ regulatory T cells ($T_{REGS}$) and total $CD8^{+}$ cells were plated in vitro for 4 days at 1:8 ratio in the presence of CD3 (1µ/mL, clone OKT-3, eBioscience) and CD28 (2µg/mL, clone CD28.2, eBioscience).

In the M-MDSCs suppression assay, sorted M-MDSCs and total $CD8^{+}$ cells were plated in vitro for 4 days at 1:2 ratio in the presence of CD3 (1µ/mL, clone OKT-3, eBioscience) and CD28 (2µg/mL, clone CD28.2, eBioscience). Cells were stimulated in vitro with Brefeldin A during the last 4 hours of the assay. Intracellular IFN$\gamma$ production and Ki-67 proliferation were measured in both assays through intracellular spectral flow cytometry.

**Gene-expression profiling analysis**

The total RNA concentration was assessed using a Qubit RNA High Sensitivity Assay (Invitrogen, Life Technologies, Waltham, MA, USA). Once the sample concentration was determined, the integrity of the total RNA was assessed using the Agilent 2100 Bioanalyzer Pico Assay (Agilent Technologies, Santa Clara, CA, USA). Samples were selected for target amplification with the GeneChip Whole Transcript (WT) Pico kit (ThermoFisher Scientific, Waltham, MA, USA).

As previously described,[5] proper RNA input was used to process the samples for whole-transcriptome expression analysis with the GeneChip WT Pico assay (ThermoFisher Scientifc). The samples were reverse-transcribed to generate amplified, fragmented, and biotinylated sense-strand cDNA (sscDNA), according to manufacturer's standard protocol. The fragmented and labeled sscDNA was then hybridized to the mouse Affimetrix Human Clariom D pico array, washed and stained using the ThermoFisher proprietary reagents in the GeneChip Fluidics Station 450 (FS450), and scanned at the GeneChip Scanner 3000 7G (ThermoFisher Scientific).

Generated CEL files were uploaded onto the Expression Console software for analysis with SST-RMA algorithm (ThermoFisher).

The standard expression microarray analysis consisted of normalizing the signal intensity distributions of all probe features on all arrays. A probe feature is a location on the array that contains many copies of the same 25-mer DNA sequence. Normalizing these distributions enabled the comparison of probe signals between groups. Next, the signal intensities for all probes in a probe set that defined a gene or exon were aggregated into a single value for each array or sample. The aggregate signal value was used to compare gene-level or exon-level expression changes between sample groups or conditions. The SST-RMA analysis algorithm incorporated pre-processing steps into the CEL files before normalization and summarization with RMA. This algorithm reduced fold change compression by applying a GC correction and by transforming the microarray data signal to a similar signal space of other methods such as RT- PCR or RNA-Seq. Differential expressed mRNAs in a comparative analysis were further identified by analysis of variance(eBay) with a p-value less than 0.05 and a fold change more than 1.1 or less than -1.1.

**Spectral flow cytometry**

Fresh PBMCs from CLL patients were seeded at 3 x $10^6$ cells/mL in culture medium and treated with CD19-4-1BBL (10 $\mu$g/ml, Roche) for 48 h. They were stimulated in vitro with Brefeldin A during the last 4 hours and stained with the surface antibodies described in Supplementary Table S5. BD Horizon Brilliant Stain Buffer (BD Biosciences) was added to the mixture of fluorescent antibodies to avoid staining artifacts. The IntraPrep Permeabilization Kit (Beckman Coulter) was used for the intracellular detection of the cytokines/transcription factors described in Supplementary Table S5. The True-Nuclear transcription factor buffer set (BioLegend) was used for the detection of Ki-67 and IFN$\gamma$ in the T-cell suppression assays described above and in Table S5. The Zombie UV Fixable Viability dye (BioLegend) staining was first performed to allow the discrimination of Live/Dead cells. Samples were acquired using an Cytek Aurora and analyzed with FCS Express 6 Flow Cytometry software (De Novo Software, Glendale, CA, USA). Fluorescence minus controls were used to determine the correct gating strategies.

**In vitro primary cell cultures for TCR sequencing**

For TCR sequencing studies, fresh PBMCs from untreated CLL patients were seeded at 3 x $10^6$ cells/mL in culture medium and treated with CD19-4-1BBL (10 µg/mlL, $\alpha$CD20 obinotuzumab (10 µg/mL), CD19-4-1BBL + $\alpha$CD20 obinotuzumab (10 µg/mL) or left untreated for 48h. Fluorescence-activated cell sorting was performed to separate CD8+ $T_{EM}$, as described above.

**RNA TCR sequencing**

RNA extraction from sorted CD8+ $T_{EM}$ was performed using Mini Kit (QIAGEN, Hilden, Germany). RNA sample quality and quantity checks (QC) were performed using an Agilent RNA 6000 Pico Assay (Agilent Technologies, Santa Clara, CA, USA). The Takara SMARTer Human TCR a/b Profiling Kit v2 has been adopted in the Cancer Genomics Laboratory at MD Anderson Cancer Center to prepare the dual indexed library. The SMARTer Human TCR a/b Profiling Kit v2 leverages SMART technology (Switching Mechanism at 5' End of RNA Template) and employs a 5' RACE-like approach to capture complete V(D)J variable regions of TCR transcripts while UMI (Unique Molecular Identifiers) is incorporated to facilitate PCR error correction and clonotype quantification at the step of data analysis.

The sequencing was performed on MiSeq using the 600-cycle MiSeq Reagent kit v3 with paired end, 2 x 300 bp reads for full-length analysis in order to cover UMI sequence, the 5' end of TRA/B V(D)J and the whole CDR3 region.

The data analysis was performed in Linux using Takara developed software Cogent NGS Immune Profiler to analyze fastq files for characterizing TCR clonality.

Further analysis was performed using the R package Immunarch (version 9.1).

**RNA extraction, RT-PCR amplification and Quantitative PCR**

RNA extraction from sorted myeloid and lymphoid cells was performed using a Mini Kit (QIAGEN, Hilden, Germany). RNA sample quality and quantity checks (QC) were performed using an Agilent RNA 6000 Pico Assay (Agilent Technologies, Santa Clara, CA, USA).

Reverse-transcription reaction from 50-200 ng RNA template was done using RevertAid™ H Minus First Strand cDNA Synthesis Kit (Fermentas, Thermo Fisher Scientific) according to the manufacturer's instructions. The sequences of primer pairs specific for each gene sequence (Sigma-Aldrich, St. Louis, MO, USA) were designed with Beacon Designer (Premier Biosoft International; Palo Alto, CA, USA) and NCBI-Primer Designing Tools (Supplementary Table S6).

Real-time PCR was performed using the SYBR® Green PCR Master Mix (Applied Biosystems, Foster City, CA, USA) with forward and reverse primers at a final concentration of 10µM. Each cDNA (2 µL) was used as template; 12.5 µL of 2x SYBR Green PCR Master Mix were mixed with template and primers. The total reaction volume was 25 µL. Three replicates for each cDNA sample were tested. 7900HT Fast

Real-Time PCR System (Applied Biosystems) was used, and cycling conditions were 10 min at 95°C (1 cycle); 10 min at 95°C (1 cycle); 15 s at 95°C plus 1 m at 60°C (40 cycles). Data were normalized to β-actin expression.

**Xenograft studies**

Eight-week-old MISTRG mice were i.v. transplanted (day 0) with $10x10^6$ MEC1 cells in 0.1 mL of saline through a 27-gauge needle. MEC1-transplanted MISTRG mice were injected i.v. on day 11 with patient-derived $CD8^+$ memory T cells, TREG, monocytes and M-MDSCs from patients with stable or high-risk CLL (Figure 7, panels B, D, F). For every patient sample, 2 tubes of total 70x10^6 PBMCs stained for myeloid cells and 2 tubes of total 70x10^6 PBMCs lymphoid cells underwent 2-hour myeloid or lymphoid 4-way sorting (as described in the sorting methods). To preserve the original lymphoid and myeloid immune cell ratio of each patient, after sorting, myeloid and lymphoid cells from each patient were mixed and transplanted into 8 mice (2 mice/group; groups: untreated, CD19-41BBL, $\alpha$CD20, $\alpha$CD20 + CD19-4-1BBL). Mice were i.p. injected with CD19-4-1BBL (1mg/kg) or $\alpha$CD20 (10mg/kg) as single agents or in combination setting or received control vehicle (saline) on days 12 and 19. In an independent survival experiment, patients-derived immune cells were injected on day 20, followed by treatment with CD19-4-1BBL or $\alpha$CD20 as single agents or in combination (days 21 and 28). Depending on the experiments, mice were monitored daily for survival or humanely euthanized at days 25-26.

**Murine cell preparations and flow cytometry**

Peripheral blood (PB) and femurs were collected from mice, and cells were isolated. Erythrocytes from bone marrow (BM) and PB samples were lysed by incubation in ammonium chloride solution (ACK) lysis buffer ($NH_4Cl$ 0.15 M, $KHCO_3$ 10 mM, $Na_2$/ethylenediaminetetraacetic acid 0.1 mM, pH 7.2-7.4) for 5 min at room temperature. After blocking of fragment crystallizable (Fc) receptors with Fc block (BD Biosciences, San Jose, CA, USA) for 10 minutes at room temperature, cells from PB, BM and SP were stained with the antibodies (15 min. at 4ºC) listed in Supplementary Tables S3-S4. Cells were analyzed with a BD LSRFortessa X-20 flow cytometer and data analyzed with FCS Express 6 Flow-Cytometry software. Absolute cell numbers were obtained by multiplying the percentage of the cells by the total number of BM cells flushed from 1 femur and tibiae.

**Histopathology and immunoistochemistry**

Collected tissues were fixed in 4% formalin for 12 h, then embedded in paraffin and five-micron thick sections were cut. For immunohistochemical (IHC) studies sections were de-paraffinized and rehydrated. IHC staining was performed on the Leica RX Bond and Leica RXm Bond using the Bond Refine DAB Detection Kit DS9800 (Leica, Nußloch, Germany). The residual wax was removed using Bond Dewax AR9222 and antibodies were diluted in Bond Diluent AR9352. Antigen retrieval reagents were Bond ER1 (Citrate Buffer) AR9961 and Bond ER2 (EDTA) AR9640. As slide washing solution was used the Bond Wash Solution 10x Concentration AR9590 which was diluted to 10% in deionized water. After the staining completion, coverslips were added and the Pathologist on site performed quality control. 12.4.6 Aperio Images scope software was used for the images.

**Statistical analysis**

The statistical analysis of the data was performed using GraphPad Prism 9.0 Software. Data were expressed as means ± standard deviations (SDs), and comparison curves or differences between experimental groups were assessed with an unpaired, 2-tailed Student *t* test (95% confidence interval) and considered statistically significant for *P*-value less than 0.05. The comparison of survival curves was performed using the log-rank test. In spectral flow experiments that involved different human myeloid cell types, normality was tested with a Shapiro–Wilk normality test; we then applied the *t* test to normally distributed data, otherwise the Mann–Whitney–Wilcoxon test. Gene set enrichment analysis was performed using the gsea() function from phenoTest package in R (Planet, 2013). The function is used to compute the enrichment scores and simulated enrichment scores for each variable and signature. For our analysis, the logscale variable was set to false, as the log2 transformed expression values were fed into the function and 20,000 simulations were used (B = 20,000). The Database for Annotation, Visualization and Integrated Discovery (DAVID) functional annotation tool was used for gene ontology and pathway (KEGG and Reactome) analysis on the list of differentially expressed genes (FDR ≤ 0.05, Log2FC more or less than 1.5/-1.5). Important GO terms and pathways were selected based on an FDR ≤ 0.05.

The outliers at end points were never excluded. The numbers of biological and technical replicates for each experiment are detailed in the figure legends. All the experiments were repeated independently and are described in the figures and figure legends. The investigators were not blinded when assessing the analyses of the experimental outcomes.

##### Graphical Representation

Heatmaps displaying significantly differentially expressed genes between groups of interest were generated in R using the function “heatmap2” from the “gplots” library. Pathways and phenotypes enriched in genes that were statistically differentially expressed were generated in R as bar plots using the “RcolorBrewer” library to display the strengths of the enrichment.

#### TABLES S1-S6

##### Supplementary Table S1. Clinical and biological features of CLL patients from MD Anderson Cancer Center

| Patient no. | Notes | Age, years | Sex | WBC | Treatment | Clinical course | CLL diagnosis (year) | ZAP70 % # | IGHV MS | FISH | Dataset |
|---|---|---|---|---|---|---|---|---|---|---|---|
| 366 | | 65 | M | 27 | Unt | progressive | 2017 | NEG | U | Trisomy12 | Clariom/RT-PCR |
| 148 | relapse | 78 | M | 109.7 | PRIOR RX | progressive | 2006 | POS | U | Del(11q) | Clariom/RT-PCR |
| 801 | | 72 | M | 74.5 | Unt | stable | 2013 | NEG | M | Trisomy12 | Clariom/RT-PCR |
| 157 | | 65 | M | 33 | Unt | stable | 2017 | POS | U | Del(13q) Del(11q) | Clariom/RT-PCR |
| 262 | | 73 | M | 87.6 | Unt | progressive | 2015 | N/A | M | Trisomy12 | TCR seq |
| 143 | | 68 | F | 61.8 | Unt | stable | 2010 | N/A | M | Del(13q) | TCR seq |
| 806 | relapse | 65 | F | 103 | PRIOR RX | progressive | 2011 | N/A | U | Trisomy12Del(17p) | TCR seq |
| 673 | | 50 | M | 137.3 | Unt | progressive | 2019 | NEG | U | Del(11q) | RT-PCR |
| 563 | | 82 | F | 158.9 | Unt | N/A | 2011 | POS | U | Del(13q) Del(11q) | Spectral Flow |
| 835 | | 76 | M | 29.4 | PRIOR RX | stable | 1991 | NEG | M | Trisomy12 | Spectral Flow |
| 276 | | 68 | F | 46.9 | Unt | stable | 2011 | NEG | M | Del(13q) | Spectral Flow |
| 715 | | 73 | F | 99.2 | Unt | stable | 2014 | POS | N/A | Del(13q) | Spectral Flow |
| 143 | | 68 | F | 61.8 | Unt | stable | 2010 | N/A | M | Del(13q) | Spectral Flow |
| 262 | | 73 | M | 87.6 | Unt | progressive | 2015 | POS | M | Trisomy12 | Spectral Flow |
| 683 | | 57 | M | 91.6 | Unt | N/A | 2017 | NEG | U | Trisomy12Del(17p) | Spectral Flow |
| 938 | | 74 | M | 77.7 | Unt | N/A | 2012 | NEG | M | Del(13q) Del(17p) | Spectral Flow |
| 457 | | 81 | F | 24.1 | PRIOR RX | stable | 2007 | N/A | M | Del(13q) | Spectral Flow |
| 547 | | 64 | F | 41.3 | Unt | progressive | 2010 | NEG | U | Trisomy12 | Spectral Flow |
| 823 | | 63 | F | 64.2 | Unt | stable | 2019 | N/A | M | Del(13q) | T-cell suppression assay |
| 176 | | 66 | M | 42.8 | Unt | stable | 2012 | NEG | M | no abnormalities | T-cell suppression assay |
| 270 | | 68 | F | 96.4 | Unt | stable | 2016 | N/A | M | N/A | T-cell suppression assay |

Abbreviations and notes: NEG, negative; POS, positive; Del, deletion; WBC, white blood cell count; IGHV MS, immunoglobulin heavy chain variable mutation status (M, mutated; U: unmutated); FISH, fluorescence *in situ* hybridization; PRIOR RX, prior therapies; Unt, untreated; N/A, not available; # Determined by flow cytometry; Dataset: dataset-related research with indicated patient sample.

#### Supplementary Table S2. Clinical and biological features of CLL patients from MD Anderson Cancer Center

| Patient no. | Notes | Age, years | Sex | WBC | Treatment | Clinical course | CLL diagnosis (year) | ZAP70 % # | IGHV MS | FISH | Dataset |
|---|---|---|---|---|---|---|---|---|---|---|---|
| *801 | | 70 | M | 57 | Unt | stable | 2011 | NEG | M | Trisomy12 | Cell depletion |
| 664 | | 70 | M | 21.1 | Unt | stable | 2009 | NEG | N/A | Del(13q) Del(17p) | Cell depletion |
| 220 | | 71 | M | 28 | PRIOR RX | stable | 2002 | NEG | M | Del(13q) | Cell depletion |
| 548 | | 62 | F | 50.6 | Unt | progressive | 2011 | NEG | M | Del(13q) | Cell depletion |
| 829 | | 61 | M | 173.9 | PRIOR RX | progressive | 2003 | NEG | M | Trisomy12 | Cell depletion |
| 253 | | 65 | M | 41.2 | Unt | progressive | 2012 | POS | U | Del(13q) | Cell depletion |
| 181 | relapse | 71 | F | 47.1 | PRIOR RX | N/A | 2010 | N/A | M | Del(11q) | Cell depletion |
| 602 | relapse | 70 | F | 31.3 | PRIOR RX | stable | 2001 | N/A | M | N/A | Cell depletion |
| 578 | | 67 | F | 26.7 | Unt | stable | 2007 | NEG | U | Del(13q) | Cell depletion |
| 419 | | 56 | F | 66.6 | Unt | stable | 2016 | NEG | M | Del(13q) | Cell depletion |
| *801 | | 70 | M | 50.9 | Unt | stable | 2011 | NEG | M | Trisomy12 | Cell depletion |
| 907 | | 63 | M | 135.5 | Unt | N/A | 2015 | N/A | U | Del(11q) | Cell depletion |
| 427 | | 65 | M | 194.1 | Unt | progressive | N/A | NEG | M | Del(13q) | Cell depletion |
| 540 | | 65 | M | 109.1 | Unt | stable | 2014 | POS | U | Del(11q) | Cell depletion |
| 665 | | 62 | M | 30 | PRIOR RX | stable | 2003 | NEG | M | N/A | Cell depletion |
| 887 | | 78 | M | 47.6 | PRIOR RX | N/A | 2012 | NEG | M | Trisomy12 Del(13q) | Cell depletion |
| 667 | | 76 | M | 222.4 | Unt | N/A | 2007 | N/A | M | Del(13q) | Cell depletion |
| 396 | | 54 | M | 70.5 | Unt | N/A | 2016 | NEG | M | N/A | Cell depletion |
| 104 | | 59 | M | 78.2 | Unt | stable | 2016 | NEG | M | N/A | Cell depletion |
| 615 | | 75 | M | 23.3 | PRIOR RX | stable | 1997 | POS | M | Del(13q) | Cell depletion |
| 070 | | 58 | M | 172 | Unt | N/A | 2013 | N/A | U | Del(11q) | Cell depletion |
| 540 | | 65 | M | 117.7 | Unt | stable | 2014 | POS | U | Del(13q) Del(11q) | Cell depletion |
| 438 | | 72 | M | 40.7 | PRIOR RX | progressive | N/A | NEG | N/A | Del(11q) | Cell depletion |
| 186 | | 56 | M | 81.9 | Unt | N/A | 2018 | POS | M | Trisomy12 Del(13q) | Cell depletion |
| 483 | | 72 | F | 30.0 | Unt | N/A | 2010 | POS | U | Trisomy12 | Cell depletion |

Abbreviations and notes: NEG, negative; POS, positive; Del, deletion; WBC, white blood cell count; IGHV MS, immunoglobulin heavy chain variable mutation status (M, mutated; U: unmutated); FISH, fluorescence *in situ* hybridization; PRIOR RX, prior therapies; Unt, untreated; N/A, not available; # Determined by flow cytometry; Dataset: dataset-related research with indicated patient sample; * different time points of the same Pt

**Supplementary Table S3. List of anti-human antibodies used for flow cytometry, myeloid cell panel**

| Antibody | Fluorochrome | Clone | Company |
|---|---|---|---|
| hCD16 | BUV 737 | 3G8 | BD Biosciences |
| hCD14 | Brilliant Violet 786 | MφP9 | BD Biosciences |
| hPDL1 | Brilliant Violet 711 | 29E-2A3 | BioLegend |
| hBCL2 | FITC | REA872 | Miltenyl Biotec |
| hHLA DR | APC-CY7 | L243 | BioLegend |
| hCD68 | PE | Y1/82A | Biolegend |
| Lineage Cocktail (hCD3/hCD19/hCD20/hCD56) | APC | UCHTI/HIB19, 2H7/5.1H11 | BioLegend |
| hCD66b | AlexaFluor 700 | G10F5 | BioLegend |

**Supplementary Table S4. List of anti-human antibodies used for flow cytometry and spectral flow cytometry, lymphoid cells**

| Antibody | Fluorochrome | Clone | Company |
|---|---|---|---|
| ***Flow cytometry:*** | | | |
| hCD8a | eFluor 450 | SK1 | eBioscience |
| hCD45RO | BUV 395 | UCHL1 | BD Biosciences |
| hCD45RA | PE-Cy7 | L48 | BD Biosciences |
| hROR1 | PE-Cy7 | 2A2 | BioLegend |
| hCD62L | Brilliant Violet 605 | DREG-56 | BioLegend |
| hCD4 | PE-Dazzle 594 | SK3 | BioLegend |
| hCD25 | PE | 4E3 | Miltenyi Biotec |
| hFoxp3 | APC | 3G3 | Miltenyi Biotec |
| hCD19 | PE-Cy5.5 | J3-119 | Beckman Coulter |
| hPDL1 | Alexa Fluor 700 | 130021 | R&D Systems |
| hCD5 | APC-Alexa 750 | BL1a | Beckman Coulter |
| hPD1 | Brilliant Violet 650 | EH12 | Biolegend |
| hBCL2 | FITC | REA872 | Miltenyl Biotec |
| ***Spectral flow cytometry:*** | | | |
| hCD4 | PE-Dazzle 594 | SK3 | BioLegend |
| hCD25 | APC/Fire810 | M-A251 | BioLegend |
| hFoxp3 | APC | 3G3 | Miltenyi Biotec |
| hCD8a | BV570 | RPA-T8 | BioLegend |
| hCD45RO | BUV 395 | UCHL | BD Biosciences |
| hCD45RA | BUV496 | HI100 | BD Biosciences |
| hCD62L | BUV563 | DREG-56 | BD Biosciences |
| hIFNy | BV711 | 4S.B3 | BD Biosciences |
| Ki-67 | BV750 | Ki67 | BioLegend |
| hIL23A | PerCP-eF710 | 23DCDP | ThermoFisher |
| TNFSF11/RANKL | PE | MIH24 | BioLegend |

#### Supplementary Table S5. List of anti-human antibodies used for Spectral flow cytometry, myeloid cell panel

| Antibody | Fluorochrome | Clone | Company |
|---|---|---|---|
| L/D | UV450 | | BioLegend |
| CD66b | PE/Fire640 | 6/40C | BioLegend |
| hCD14 | BV785 | 63DE | Biolegend |
| hCD16 | BUV737 | 3G8 | BD |
| hHLA DR | APC-CY7 | L243 | BioLegend |
| hCD68 | PE | Y1/82A | Biolegend |
| Lineage (hCD3/hCD19/hCD20/hCD56) | BV570 | UCHTI/HIB19, 2H7/5.1H11 | BioLegend |
| MRC1 | BV605 | 15-2 | BioLegend |
| hCD163 | BV711 | GHI/61 | Biolegend |
| PD-L1 | BV650 | 29E.2A3 | BioLegend |
| IL-6 | PerCP-Cy5.5 | MQ2-13A5 | Biolegend |
| BCL-2 | PE-CF594 | BCL-2/100 | BD |
| NOTCH4 | BV421 | MHN4-2 | BD |
| hCD62L | BUV563 | SK11 | BD |
| HDAC8 | AlexaFluor 647 | E5 | Santa Cruz Biotechnology |
| PAWR | AlexaFluor 546 | A10 | Santa Cruz Biotechnology |
| 41BB | BV750 | 4B4-1 | BioLegend |
| IFNa | PACB | 7N41 | Life technologies |
| hCDKN1B | AlexaFluor 532 | | Life technologies |
| hCD24 | BUV496 | ML5 | BD |
| MCOLN2 | AlexaFluor 488 | F1 | Santa Cruz Biotechnology |
| GAS6 | AlexaFluor 680 | A-9 | Santa Cruz Biotechnology |
| EIF4E | AlexaFluor 594 | P2 | Santa Cruz Biotechnology |
| IRF4 | PE | 3E4 | Santa Cruz Biotechnology |
| hCD11c | BUV661 | B-LY6 | BD |
| hCD1a | BUV805 | SK9 | BD |
| hCD1c | BV510 | L161 | BioLegend |
| hCD80 | BUV395 | L307.4 | BD |
| hCD86 | BUV615 | BU63 | BD |

#### Supplementary Table S6. Primers for qRT-PCR analysis

| | HUMAN | |
|---|---|---|
| **GENE** | **FORWARD** | **REVERSE** |
| *RELA* | 5’-TGGGAATCCAGTGTGTGAAG-3’ | 5’-CACAGCATTCAGGTCGTAGT-3’ |
| *TAB3* | 5’-CACTATAGCCAGCGTCCTTTAC-3’ | 5’-GAAGGAGGTGGTGAATGGTAAG-3’ |
| *EIF2AK2* | 5’-GCTTCCCTTTCTCCTGTTACT-3’ | 5’-TGGGTTCCATCAGTGTTATCC-3’ |
| *TP53BP2* | 5’-GCTAAAGTACCACCTCCTGTTC-3’ | 5’-GAGAACTTCCGTCTGGCTTAAT-3’ |
| *TNFSF11* | 5’-CCCAAGTTCTCATACCCTGATG-3’ | 5’-TTCCTCTCCAGACCGTAACT-3’ |
| *IL23A* | 5’-GTGGAAGTGGGCAGAGATT-3’ | 5’-CAGCAACAGCAGCATTACAG-3’ |
| *CD62L* | 5’-CCTTCTTCAGCCACCTCTCTTT-3’ | 5’-AGCGCAGGCTATTTCTCTCTTC-3’ |
| *CDKN1B* | 5’-AACTCTGAGGACACGCATTTGG-3’ | 5’-GGTTGCAGGTCGCTTCCTTATT-3’ |
| *PAWR* | 5’-TGATGAAGCAGGGCAGAAAGA-3’ | 5’-TAGGTGGCTCCTGTAGCAGATA-3’ |
| *NOTCH4* | 5’-GGAGGATATCGATGAGTGCAGAAG-3’ | 5’-TTCAAAGCCTGGGAGACACTT-3’ |
| *MAPK3K13* | 5’-TGGAGAGTAGCAGGGAGACTAA-3’ | 5’-GCTGTTGAGCGTAACTGAAACC-3’ |
| *IFNA* | 5’-CTTGATGCTCCTGGCACAAATG-3’ | 5’-CTGGAACTGGTTGCCATCAAAC-3’ |
| *CD24* | 5’-AGTCTCTTCGTGGTCTCACTCT-3’ | 5’-TGGACTTCCAGACGCCATTT-3’ |
| *MCOLN2* | 5’-TCTTGGAACCTCTACGCTCTTG-3’ | 5’-CAGTGAGGCCTGCATTGTTAAA-3’ |
| *GAS6* | 5’-GGACCTCGTGCAGCCTATAAA-3’ | 5’-CGTGTTCACTTTCACCGTTTCC-3’ |
| *KPNA3* | 5’-TGTGATGTCCTGTCACACTTCC-3’ | 5’-CTGGTTGCCTGCTGTTATGTTG-3’ |
| *EIF4E* | 5’-GGTATTGAGCCTATGTGGGAAGATG-3’ | 5’-GGTCACTTCGTCTCTGCTGTTT-3’ |
| *IRF4* | 5’-CAGCCCAGGTTCACAACTACA-3’ | 5’-TGGGACATTGGTACGGGATTTC-3’ |
| *HDAC8* | 5’-TTTGGGAGGAGGAGGCTATAA-3’ | 5’-CTGGGATCTCAGAGGATAGTGT-3’ |

**LEGENDS FOR TABLES S7-S11**

**Table S7.**

Coding and non-coding RNAs downregulated (Down) or upregulated (Up) in Classical monocytes (C), Intermediate monocytes (INT), MDSC, Non-classical monocytes (NC), CD8$^{+}$ central memory T cells (TCM), CD8$^{+}$ effector memory T cells (TEM), CD4$^{+}$ regulatory T cell (TREG) treated with CD19-4-1BBL (BB) compared with untreated (UNTR) control cells.

**Table S8.**

Commonly downregulated or upregulated RNAs in untreated Intermediate monocytes (INT), MDSC and CD4$^{+}$ regulatory T cells.

**Table S9.**

Gene ontology analysis performed on differentially expressed genes (FDR <= 0.05, Log2FC

more or less than 1.5/-1.5) related to Classical monocytes (C), Intermediate monocytes (INT), MDSC, Non-classical monocytes (NC), CD8$^{+}$ central memory T cells (TCM), CD8$^{+}$ effector memory T cells (TEM), CD4$^{+}$ regulatory T cell (TREG) treated with CD19-4-1BBL (BB) compared with untreated (UNTR) control cells. GO terms were selected based on an FDR <= 0.05.

**Table S10.**

Pathway analysis performed on differentially expressed genes (FDR <= 0.05, Log2FC more or less than 1.5/-1.5) related to Classical monocytes (C), Intermediate monocytes (INT), MDSC, Non-classical monocytes (NC), CD8$^{+}$ central memory T cells (TCM), CD8$^{+}$ effector memory T cells (TEM), CD4$^{+}$ regulatory T cell (TREG) treated with CD19-4-1BBL (BB) compared with untreated (UNTR) control cells. Pathways were selected based on an FDR <= 0.05.

**Table S11.**

The table shows selected genes of interest (represented on supervised Heatmaps) differentially expressed on Classical monocytes (C), Intermediate monocytes (INT), MDSC, CD8$^{+}$ central memory T cells (TCM), CD8$^{+}$ effector memory T cells (TEM), CD4$^{+}$ regulatory T cell (TREG) treated with CD19-4-1BBL (BB) compared with untreated (UNTR) control cells. The genes with FDR <= 0.05, Log2FC more or less than 1.5/-1.5 have been retained as differentially expressed.

References and Pathway categories are included.

**Supplementary Table S12. TRA gene usage of $T_{EM}$ treated with CD19-4-1BBL, CD19-4-1BBL + αCD20 or left untreated.**

| | Genes | CD19-41BBL | CD19-4-1BBL + αCD20 | Untreated |
|---|---|---|---|---|
| 1 | TRAV1-1 | 27 | 37 | 23 |
| 2 | TRAV1-2 | 187 | 190 | 152 |
| 3 | TRAV10 | 43 | 56 | 39 |
| 4 | TRAV12-1 | 71 | 148 | 57 |
| 5 | TRAV12-2 | 145 | 209 | 125 |
| 6 | TRAV12-3 | 45 | 115 | 45 |
| 7 | TRAV13-1 | 118 | 187 | 105 |
| 8 | TRAV13-2 | 58 | 63 | 51 |
| 9 | TRAV14DV4 | 95 | 123 | 88 |
| 10 | TRAV16 | 38 | 28 | 31 |
| 11 | TRAV17 | 107 | 210 | 92 |
| 12 | TRAV19 | 83 | 99 | 55 |
| 13 | TRAV2 | 33 | 80 | 29 |
| 14 | TRAV20 | 50 | 77 | 48 |
| 15 | TRAV21 | 106 | 172 | 87 |
| 16 | TRAV22 | 45 | 61 | 42 |
| 17 | TRAV23DV6 | 41 | 33 | 26 |
| 18 | TRAV24 | 44 | 54 | 41 |
| 19 | TRAV25 | 27 | 63 | 37 |
| 20 | TRAV26-1 | 29 | 97 | 33 |
| 21 | TRAV26-2 | 46 | 83 | 36 |
| 22 | TRAV27 | 59 | 113 | 50 |
| 23 | TRAV28 | NA | 1 | NA |
| 24 | TRAV29DV5 | 102 | 143 | 81 |
| 25 | TRAV3 | 77 | 114 | 65 |
| 26 | TRAV30 | 20 | 39 | 11 |
| 27 | TRAV34 | 3 | 10 | 3 |
| 28 | TRAV35 | 35 | 71 | 28 |
| 29 | TRAV36DV7 | 4 | 38 | 8 |
| 30 | TRAV38-1 | 59 | 143 | 39 |
| 31 | TRAV38-2DV8 | 83 | 240 | 61 |
| 32 | TRAV39 | 11 | 18 | 7 |
| 33 | TRAV4 | 75 | 100 | 68 |
| 34 | TRAV40 | 2 | 7 | 1 |
| 35 | TRAV41 | 34 | 47 | 23 |
| 36 | TRAV5 | 91 | 116 | 64 |
| 37 | TRAV6 | 27 | 35 | 16 |
| 38 | TRAV8-1 | 51 | 78 | 32 |
| 39 | TRAV8-2 | 39 | 49 | 37 |
| 40 | TRAV8-2, TRAV8-4 | NA | NA | 1 |
| 41 | TRAV8-3 | 50 | 69 | 45 |
| 42 | TRAV8-3, TRAV8-5 | NA | 1 | NA |
| 43 | TRAV8-4 | 47 | 72 | 32 |
| 44 | TRAV8-4, TRAV8-2 | NA | 1 | NA |
| 45 | TRAV8-4, TRAV8-6 | 1 | NA | NA |
| 46 | TRAV8-5 | 3 | NA | NA |
| 47 | TRAV8-6 | 47 | 85 | 45 |
| 48 | TRAV9-1 | 3 | 2 | 1 |

| | | | | |
|---|---|---|---|---|
| 49 | TRAV9-2 | 28 | 58 | 22 |
| 50 | TRDV1 | 20 | 37 | 18 |
| 51 | TRDV3 | 2 | NA | 1 |

**Supplementary Table S13. TRB gene usage of $T_{EM}$ treated with CD19-4-1BBL, CD19-4-1BBL + αCD20 or left untreated**

| | Genes | CD19-41BBL | CD19-4-1BBL + αCD20 | Untreated |
|---|---|---|---|---|
| 1 | TRBV1 | 2 | 10 | 2 |
| 2 | TRBV10-1 | 25 | 61 | 18 |
| 3 | TRBV10-2 | 44 | 162 | 31 |
| 4 | TRBV10-3 | 63 | 395 | 49 |
| 5 | TRBV11-1 | 5 | 50 | 3 |
| 6 | TRBV11-2 | 105 | 411 | 76 |
| 7 | TRBV11-2, TRBV7-2 | *NA* | 1 | *NA* |
| 8 | TRBV11-3 | 16 | 92 | 15 |
| 9 | TRBV11-3, TRBV11-1 | 1 | 4 | 1 |
| 10 | TRBV12-1 | 1 | 5 | 1 |
| 11 | TRBV12-2 | 3 | 22 | *NA* |
| 12 | TRBV12-3 | 65 | 578 | 46 |
| 13 | TRBV12-3, TRBV12-4 | 1 | 14 | *NA* |
| 14 | TRBV12-4 | 138 | 501 | 126 |
| 15 | TRBV12-4, TRBV12-3 | *NA* | 1 | *NA* |
| 16 | TRBV12-5 | 6 | 39 | 9 |
| 17 | TRBV13 | 56 | 286 | 41 |
| 18 | TRBV13, TRBV9 | *NA* | 1 | *NA* |
| 19 | TRBV14 | 60 | 214 | 41 |
| 20 | TRBV15 | 93 | 486 | 77 |
| 21 | TRBV16 | 1 | 14 | 2 |
| 22 | TRBV18 | 40 | 261 | 26 |
| 23 | TRBV19 | 135 | 564 | 114 |
| 24 | TRBV2 | 131 | 642 | 111 |
| 25 | TRBV20-1 | 314 | 2063 | 227 |
| 26 | TRBV21-1 | 16 | 127 | 17 |
| 27 | TRBV23-1 | 16 | 21 | 17 |
| 28 | TRBV24-1 | 57 | 379 | 27 |
| 29 | TRBV25-1 | 35 | 200 | 42 |
| 30 | TRBV27 | 232 | 1263 | 206 |
| 31 | TRBV28 | 142 | 1819 | 142 |
| 32 | TRBV29-1 | 141 | 810 | 103 |
| 33 | TRBV3-1 | 83 | 520 | 77 |
| 34 | TRBV3-2 | 1 | 10 | *NA* |
| 35 | TRBV30 | 41 | 483 | 25 |
| 36 | TRBV4-1 | 117 | 619 | 89 |
| 37 | TRBV4-1, TRBV4-2 | *NA* | 1 | *NA* |
| 38 | TRBV4-1, TRBV4-3 | *NA* | 2 | *NA* |
| 39 | TRBV4-2 | 73 | 459 | 52 |
| 40 | TRBV4-2, TRBV4-3 | *NA* | 2 | *NA* |
| 41 | TRBV4-3 | 37 | 605 | 1 |
| 42 | TRBV4-3, TRBV4-2 | *NA* | 1 | *NA* |
| 43 | TRBV5-1 | 156 | 927 | 140 |
| 44 | TRBV5-1, TRBV5-8 | 1 | *NA* | *NA* |
| 45 | TRBV5-2 | *NA* | 1 | *NA* |
| 46 | TRBV5-3 | *NA* | 4 | *NA* |
| 47 | TRBV5-4 | 48 | 286 | 47 |

| | | | | |
|---|---|---|---|---|
| 48 | TRBV5-5 | 42 | 277 | 24 |
| 49 | TRBV5-6 | 70 | 453 | 48 |
| 50 | TRBV5-6, TRBV5-5 | *NA* | 2 | *NA* |
| 51 | TRBV5-7 | NA | 2 | NA |
| 52 | TRBV5-8 | 8 | 49 | 6 |
| 53 | TRBV6-1 | 204 | 587 | 162 |
| 54 | TRBV6-1, TRBV6-4 | NA | 2 | NA |
| 55 | TRBV6-1, TRBV6-4, TRBV6-5, TRBV6-6 | NA | 5 | NA |
| 56 | TRBV6-1, TRBV6-5, TRBV6-6 | NA | 1 | NA |
| 57 | TRBV6-2 | 4 | 16 | 4 |
| 58 | TRBV6-2, TRBV6-3 | 76 | 563 | 44 |
| 59 | TRBV6-2, TRBV6-3, TRBV6-4, TRBV6-5, TRBV6-6 | NA | 1 | NA |
| 60 | TRBV6-2, TRBV6-3, TRBV6-5 | NA | 1 | NA |
| 61 | TRBV6-2, TRBV6-3, TRBV6-5, TRBV6-6 | NA | 2 | NA |
| 62 | TRBV6-3 | NA | 22 | NA |
| 63 | TRBV6-3, TRBV6-2 | 26 | 100 | 8 |
| 64 | TRBV6-4 | 136 | 301 | 123 |
| 65 | TRBV6-4, TRBV6-5 | NA | 1 | NA |
| 66 | TRBV6-4, TRBV6-5, TRBV6-6 | NA | 1 | NA |
| 67 | TRBV6-5 | 100 | 565 | 82 |
| 68 | TRBV6-5, TRBV6-2, TRBV6-3 | NA | 1 | NA |
| 69 | TRBV6-5, TRBV6-6 | 1 | 10 | NA |
| 70 | TRBV6-6 | 64 | 294 | 44 |
| 71 | TRBV6-6, TRBV6-5 | NA | 2 | NA |
| 72 | TRBV6-7 | NA | 7 | 1 |
| 73 | TRBV6-9 | NA | 5 | NA |
| 74 | TRBV7-2 | 64 | 562 | 62 |
| 75 | TRBV7-2, TRBV7-8 | NA | 1 | NA |
| 76 | TRBV7-3 | 46 | 349 | 52 |
| 77 | TRBV7-3, TRBV7-6 | NA | 1 | NA |
| 78 | TRBV7-3, TRBV7-8 | NA | 1 | NA |
| 79 | TRBV7-3, TRBV7-9 | 1 | NA | NA |
| 80 | TRBV7-4 | 2 | 9 | 1 |
| 81 | TRBV7-4, TRBV7-3 | NA | 1 | NA |
| 82 | TRBV7-6 | 36 | 391 | 42 |
| 83 | TRBV7-7 | 10 | 91 | 12 |
| 84 | TRBV7-8 | 87 | 478 | 77 |
| 85 | TRBV7-9 | 243 | 1646 | 199 |
| 86 | TRBV7-9, TRBV11-1 | NA | 2 | 1 |
| 87 | TRBV9 | 139 | 723 | 105 |

**Table S14.**

Epitopes and related Tumor-associated Antigens (TAA) predicted through the VDJdb database related to the CDR3 sequences of the TRA genes identified on CD8+ effector memory T cells treated in vitro with CD19-4-1BBL and aCD20.

**Table S15.**

Epitopes and related Tumor-associated Antigens (TAA) predicted through the VDJdb database related to the CDR3 sequences of the TRB genes identified on CD8+ effector memory T cells treated in vitro with CD19-4-1BBL and aCD20.

**Table S16.**

Epitopes and related Antigens predicted through the VDJdb and McPAS-TCR databases related to the CDR3 sequences of the TRB genes identified on CD8+ effector memory T cells (Pt sample #143) before treatment.

**Table S17.**

Epitopes and related Antigens predicted through the VDJdb and McPAS-TCR databases related to the CDR3 sequences of the TRB genes identified on CD8+ effector memory T cells (Pt sample #143) after treatment with CD19-4-1BBL + aCD20.

**Table S18.**

Epitopes and related Antigens predicted through the VDJdb and McPAS-TCR databases related to the CDR3 sequences of the TRB genes identified on CD8+ effector memory T cells (Pt sample #262) before treatment.

**Table S19.**

Epitopes and related Antigens predicted through the VDJdb and McPAS-TCR databases related to the CDR3 sequences of the TRB genes identified on CD8+ effector memory T cells (Pt sample #262) after treatment with CD19-4-1BBL + aCD20.

**Table S20.**

Epitopes and related Antigens predicted through the VDJdb and McPAS-TCR databases related to the CDR3 sequences of the TRB genes identified on CD8+ effector memory T cells (Pt sample #806) before treatment.

**Table S21.**

Epitopes and related Antigens predicted through the VDJdb and McPAS-TCR databases related to the CDR3 sequences of the TRB genes identified on CD8+ effector memory T cells (Pt sample #806) after treatment with CD19-4-1BBL + aCD20.

#### Supplementary Table S22. Clinical and biological features of CLL patient samples of survival in vivo studies

| Patient no. | CLL diagnosis (y) | Rai stage | Infection history | WBC | PRIOR RX | ZAP70 % # | IGHV | Gene mutation | FISH | Survival | Survival (GA101) | Survival (CD19-4-1BBL) | Survival (GA101 + CD19-41BBL) |
|---|---|---|---|---|---|---|---|---|---|---|---|---|---|
| 160 | 2014 | N/A | N | 115 | U | + | M | NO | Del(13q) | 33 | 33 | 24 | 88 |
| 160 | 2014 | N/A | N | 115 | U | + | M | NO | Del(13q) | 48 | 66 | 44 | |
| 160 | 2014 | N/A | N | 115 | U | + | M | NO | Del(13q) | | | 44 | |
| 782 | 2011 | N/A | N | 31.6 | U | - | U | NO | Del(13q) | 29 | 29 | 29 | 33 |
| 782 | 2011 | N/A | N | 31.6 | U | - | U | NO | Del(13q) | 29 | 73 | 36 | 66 |
| 958 | 2001 | I | Y | 40.9 | U | - | U | NO | NEG | 29 | 36 | 30 | 32 |
| 958 | 2001 | I | Y | 40.9 | U | - | U | NO | NEG | 53 | 68 | | 53 |
| 958 | 2001 | I | Y | 40.9 | U | - | U | NO | NEG | | | | 92 |
| 036 | 2020 | N/A | Y | 292.9 | U | N/A | M | NO | Del(13q) | 23 | 25 | 22 | 43 |
| 036 | 2020 | N/A | Y | 292.9 | U | N/A | M | NO | Del(13q) | 48 | 63 | 27 | 60 |
| 408 | 2009 | N/A | Y | 94.9 | U | N/A | M | BIRC3 | Trisomy12 | 22 | 22 | 27 | 72 |
| 408 | 2009 | N/A | Y | 94.9 | U | N/A | M | BIRC3 | Trisomy12 | 42 | 72 | 28 | 329 |
| 555 | 2019 | N/A | Y | 168.4 | U | N/A | U | TP53 | Del(13q)<br>Del(11q) | 29 | 24 | 24 | 57 |
| 555 | 2019 | N/A | Y | 168.4 | U | N/A | U | TP53 | Del(13q)<br>Del(11q) | 42 | 24 | 38 | 63 |
| 357 | 1999 | N/A | N/A | 120.5 | Y | N/A | M | TP53 | Del(13q) | 25 | 28 | 24 | 24 |
| 357 | 1999 | N/A | N/A | 120.5 | Y | N/A | M | TP53 | Del(13q) | 27 | | 44 | 357 |
| 624 | 2014 | N/A | Y | 93.6 | Y | N/A | U | | NEG | 25 | 24 | 34 | 25 |
| 624 | 2014 | N/A | Y | 93.6 | Y | N/A | U | | NEG | 25 | 25 | | 34 |
| 806 | 2011 | IV | Y | 95.9 | Y | N/A | U | TP53<br>SF3B1<br>BIRC3 | Del(17p)<br>Trisomy12 | 25 | 24 | 25 | 37 |
| 806 | 2011 | IV | Y | 95.9 | Y | N/A | U | TP53<br>SF3B1<br>BIRC3 | Del(17p)<br>Trisomy12 | | | 25 | |

Abbreviations and notes: Del, deletion; WBC, white blood cell count; IGHV MS, immunoglobulin heavy chain variable mutation status (M, mutated; U: unmutated); Gene mutation: NGS-based analysis for the detection of somatic mutations performed on DNA; FISH, fluorescence *in situ* hybridization; PRIOR RX, prior therapies; Unt, untreated; N/A, not available; # Determined by flow cytometry; Survival: survival time (days) upon indicated treatment described in Figure 7. Immune cells from samples 357, 624, 806 indicated in green have been injected into mice starting at day 20.

### Figure S1 related to Figure 1

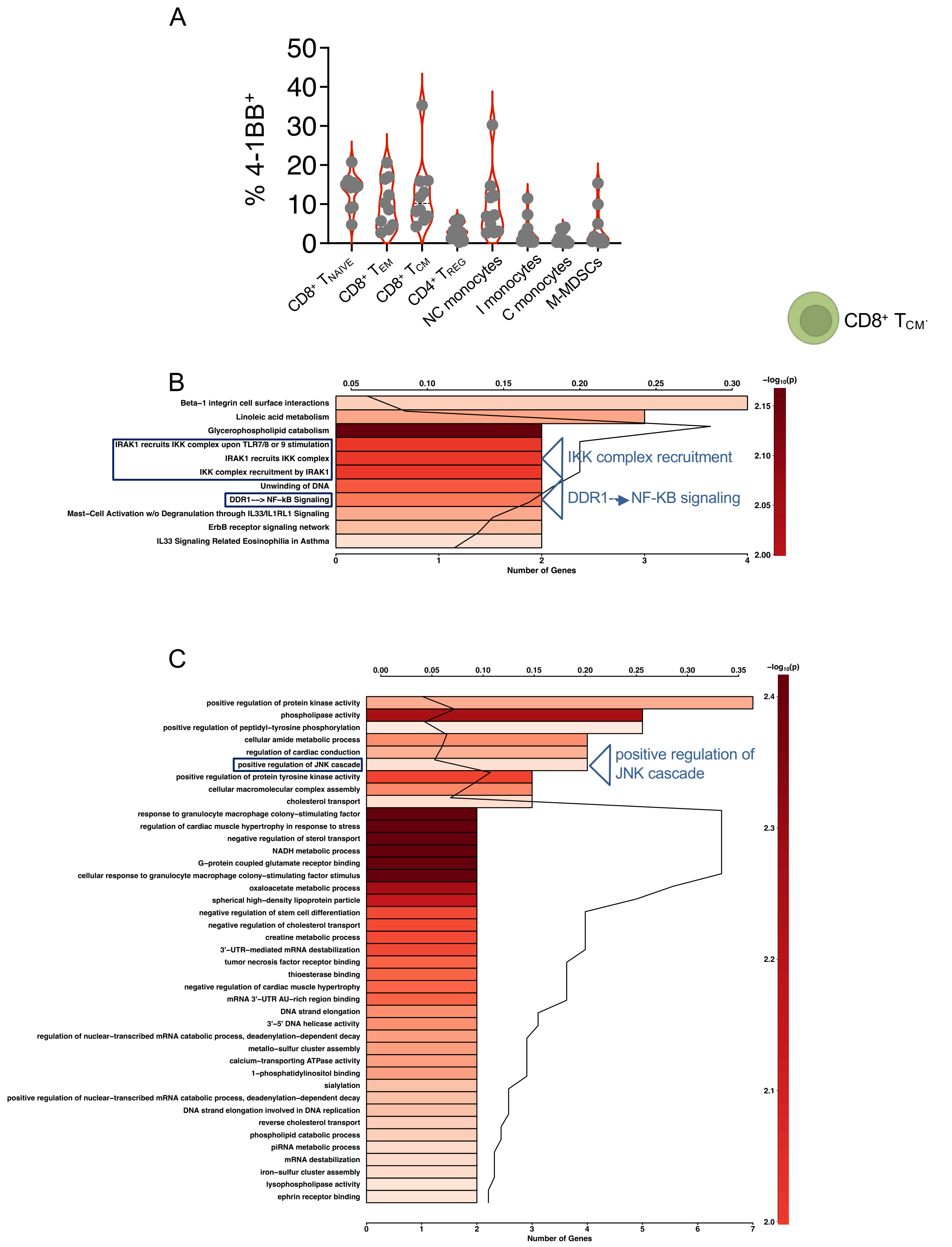

**Fig. S1. Transcriptome analysis of patient-derived central memory $CD8^+$ T cells exposed to CD19-4-1BBL.** (A) The relative contribution of $CD8^+$ $CD45RO^-CD45RA^+$ $CD62L^+$ Naïve, $CD8^+$ $CD45RA^-$ $CD45RO^+$ $CD62L^+$ $T_{CM}$, $CD4^+$ $CD25^+$ $FoxP3^+$ $T_{REG}$, $CD14^+CD16^{++}$ NC, $CD14^{++}$ $CD16^+$ I, $CD14^{++}CD16^-$ monocytes, $CD14^+HLA\text{-}DR^{low/-}$ M-MDSCs expressing 4-1BB, from PBMCs of patients with CLL (n=10, Table S1) is shown. (B-C) Fresh PBMCs from patients with CLL (n = 4, Table S1) were incubated with CD19-4-1BBL (10μ/mL) for 22 h. RNA was isolated from FACS sorted cells and a human Clariom D Pico assay was used to perform a broad transcriptome gene- and exon- level analysis of coding RNA. (B) Pathway and (C) GO analysis of differentially expressed transcripts between CD19-41BBL treated and untreated $CD8^+$ $T_{CM}$ are shown.

### Figure S2 related to Figure 2

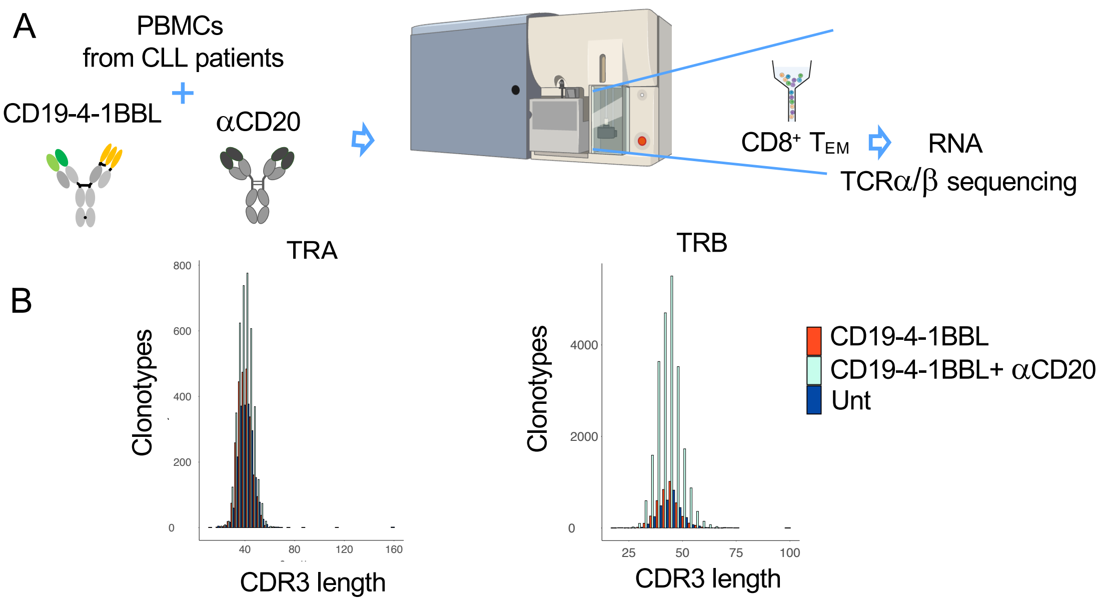

**Fig. S2. TCR sequencing of effector memory CD8$^+$ T cells upon CD19-4-1BBL treatment.** (A) Fresh PBMCs from patients with CLL (n = 3, Table S1) and age-matched healthy donor control (n = 3) were incubated with CD19-4-1BBL (10μ/ml), CD19-4-1BBL + $\alpha$CD20 obinotuzumab (10 μg/mL) or left untreated for 48 h. RNA was isolated from FACS sorted hCD8$^+$ CD45RA$^-$CD45RO$^+$CD62L$^-$ $T_{EM}$ and the Takara SMARTer Human TCR a/b Profiling Kit v2 has been performed. (B) The graph shows the CDR3 length distribution of TRA and TRB genes related to the CD8$^+$ $T_{EM}$ treated with CD19-4-1BBL, CD19-4-1BBL + $\alpha$CD20 obinotuzumab or left untreated.

### Figure S3 related to Figure 2

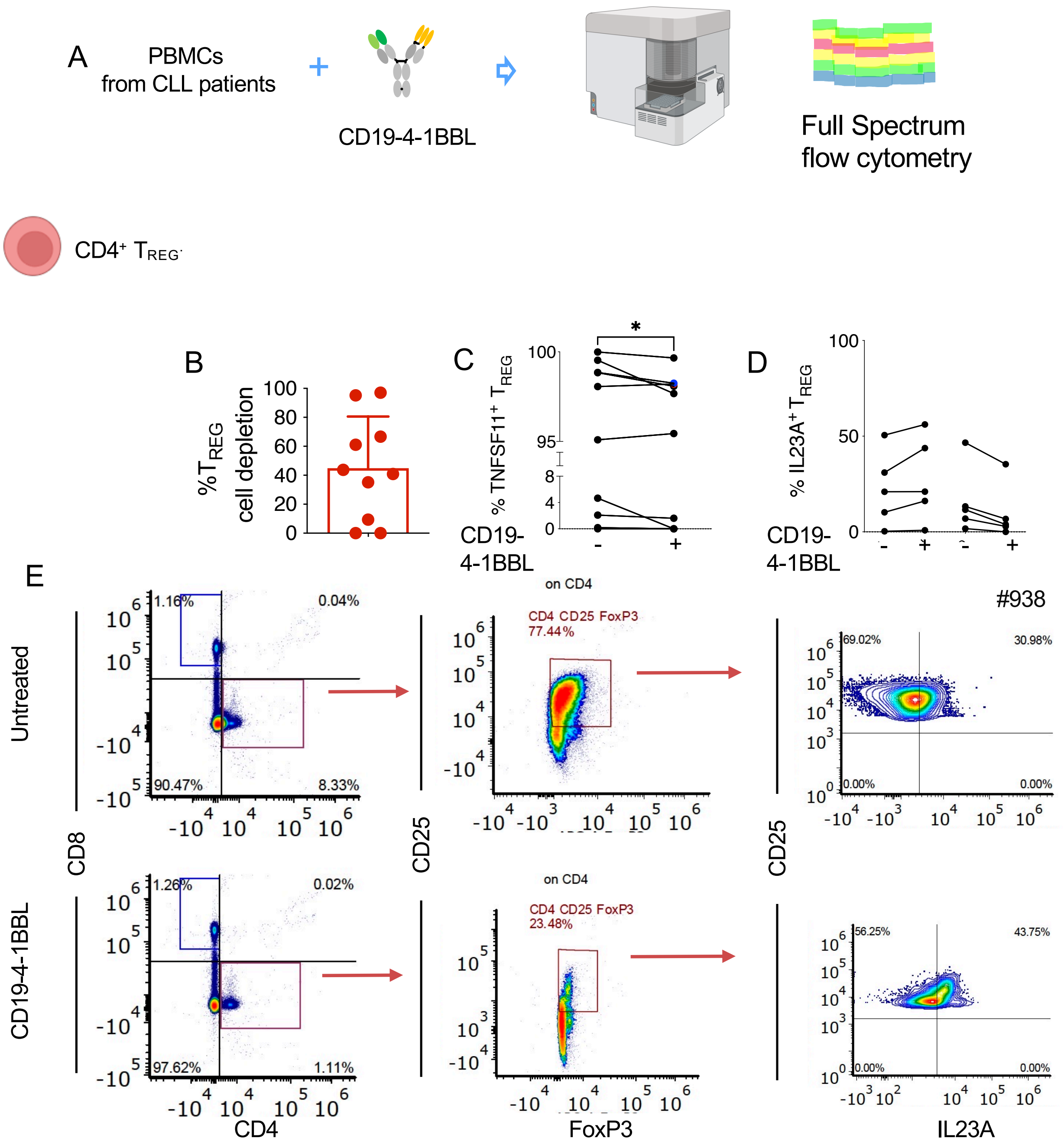

**Fig. S3. Costimulatory activity of CD19-4-1BBL in patient-derived CD4$^+$ $T_{REG}$ cells**

(A) Fresh PBMCs from CLL patients (n = 10) were incubated with CD19-4-1BBL (10 $\mu$g/mL, Roche) for 48 h and then stained and analyzed by spectral flow cytometry. (B) Regulatory hCD4$^+$CD25$^+$Foxp3$^+$ $T_{REG}$ cell depletion (n = 10, Supplementary Table S1) after 48 h of treatment with 10µg/mL of CD19-4-1BBL was analyzed by spectral flow cytometry and calculated by the formula: the following formula: 100 - % remaining cells, where % remaining cells = (absolute number in treated samples/absolute number in untreated samples) × 100. Details are described in the Supplementary Methods. (C) the relative contribution of hCD4$^+$CD25$^+$Foxp3$^+$ TNFS11$^+$ $T_{REG}$ cells treated with CD19-4-1BBL or left untreated is shown. *P* value is given by Mann-Whitney-Wilcoxon test *$P < 0.05$. (D) the relative contribution of hCD4$^+$CD25$^+$Foxp3$^+$ IL23$^+$ $T_{REG}$ cells treated with CD19-4-1BBL or left untreated and (E) related dot plots from representative patient #938 are shown.

### Figure S4 related to Figure 4

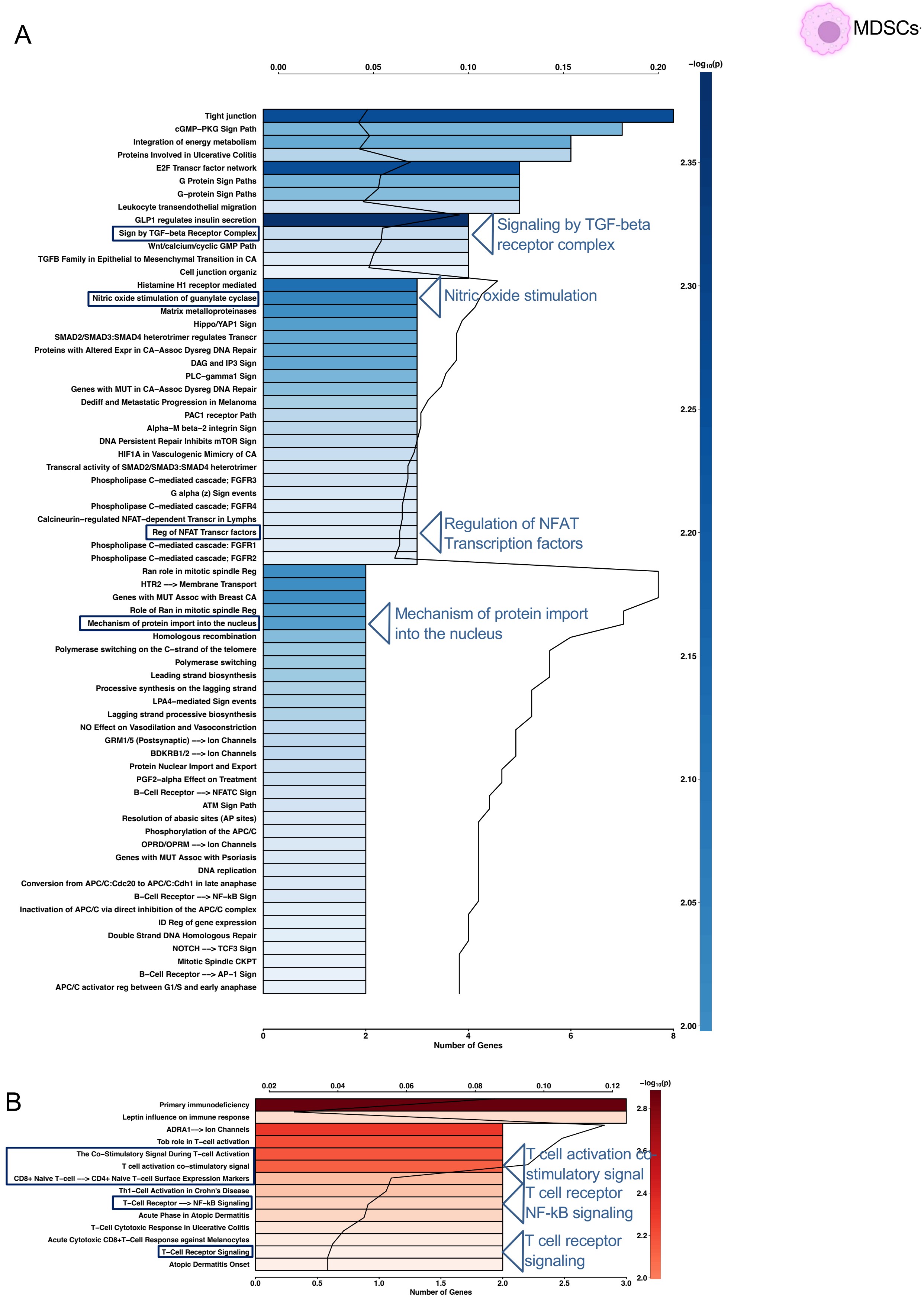

**Fig. S4. Transcriptome analysis of patient-derived monocytic myeloid-derived suppressor cells**

(A) GO and (B) pathway analysis of differentially expressed transcripts between CD19-41BBL treated and untreated M-MDSCs are shown.

### Figure S5 to Figure 4

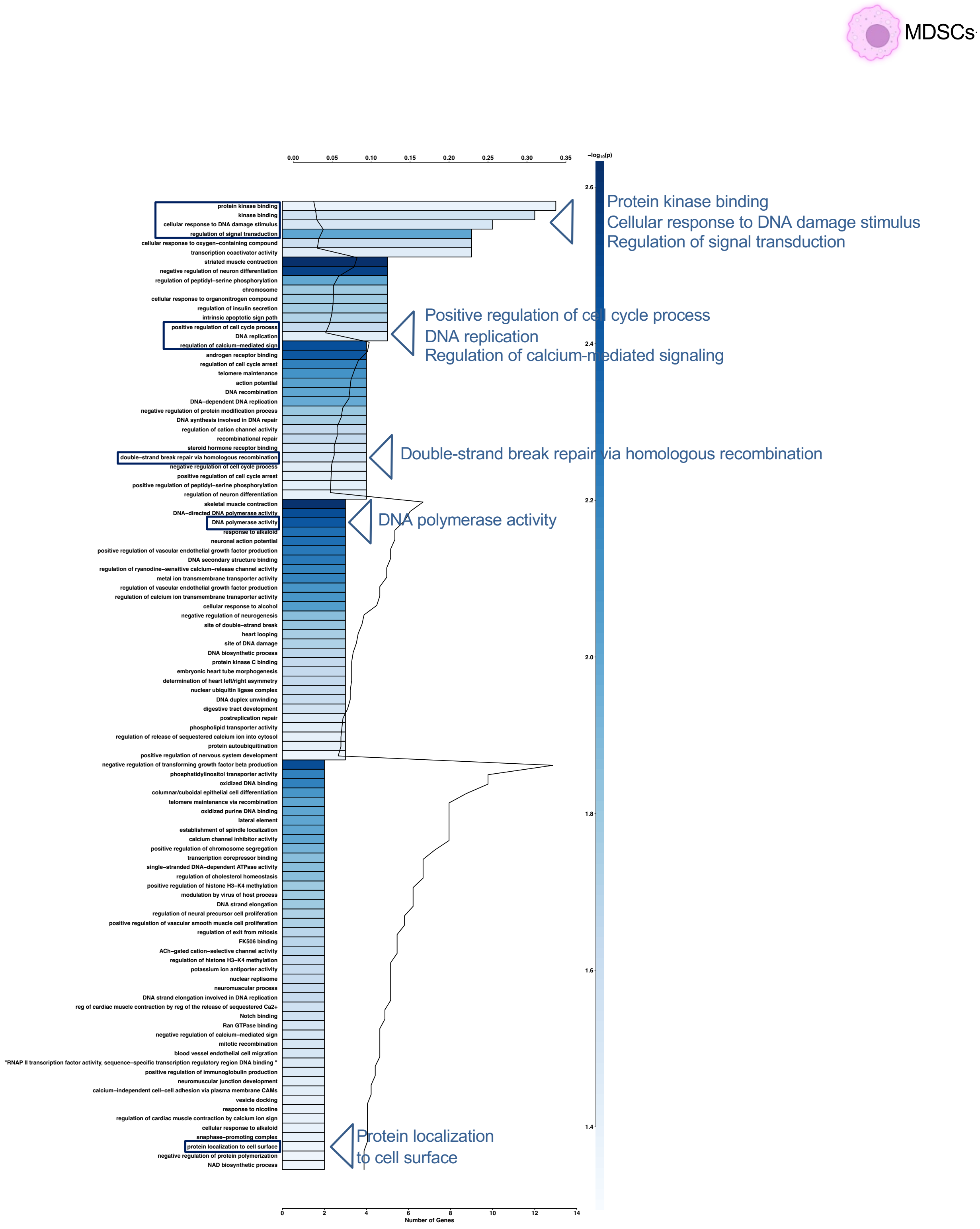

**Fig. S5. Transcriptome analysis of patient-derived monocytic myeloid-derived suppressor cells**

(A) GO analysis of differentially expressed transcripts between CD19-41BBL treated and untreated M-MDSCs is shown.

### Figure S6 related to Figure 7

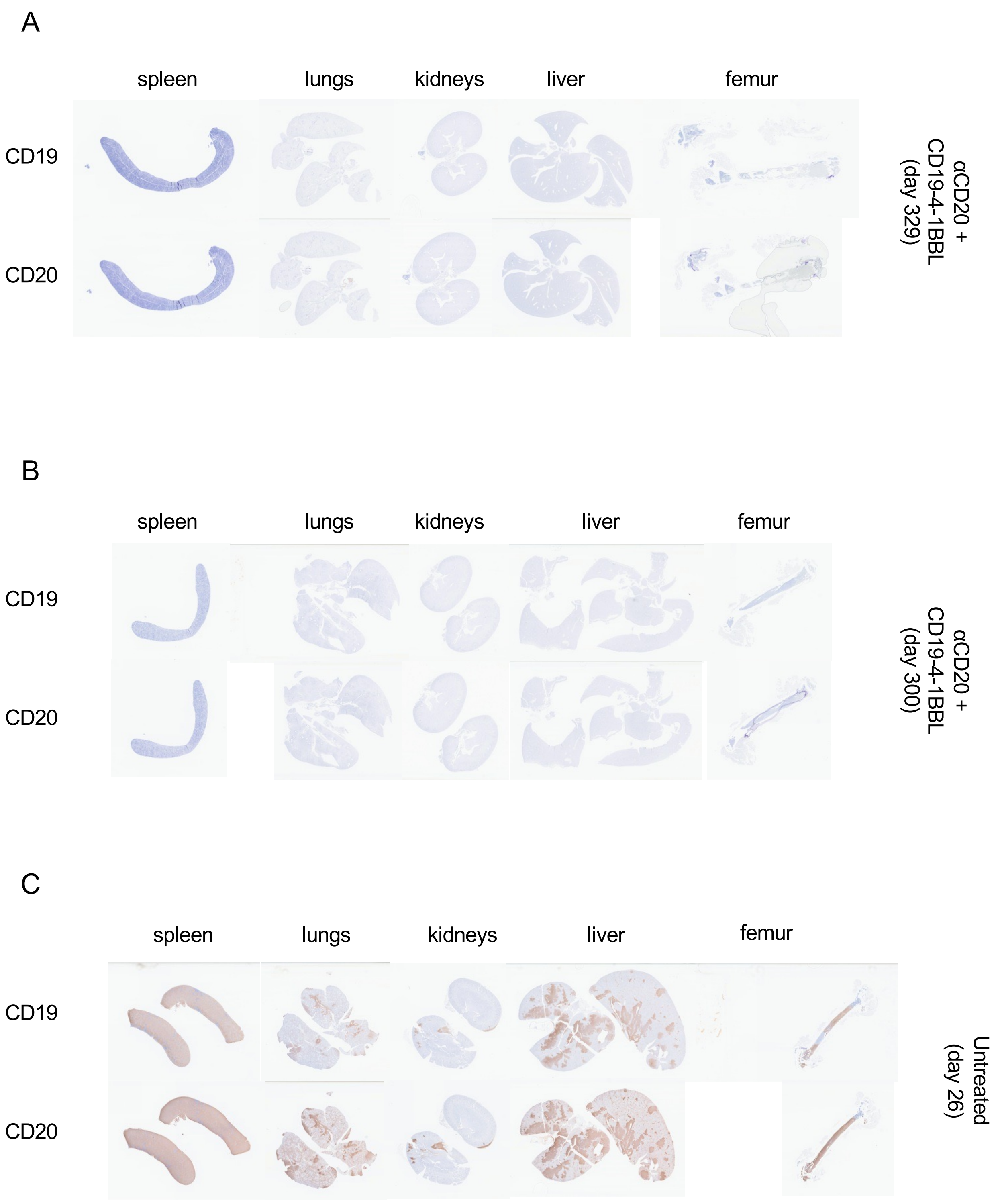

**Figure S6. Histopathological analysis of long survivor xenotransplanted MISTRG mice.**

(A-B) Immunohistochemical (IHC) stains for anti-human CD19 and CD20 of spleen, lungs, kidneys, livers and femurs of representative MISTRG mice transplanted intravenously (i.v.) with MEC1 cells (day 0), that received weekly injections of CD19-4-1BBL (1mg/kg) and $\alpha$CD20 (10mg/kg) combination, after the adoptive transfer of patient-derived immune cells (days 11 or day 20, depending on the experiment). (A) IHC stains related to the representative mouse killed at day 329, described in Figure 7E. (B) IHC stains related to representative mouse that was found dead at day 300, described in Figure 7G. (C) IHC stains related to a representative MEC1-transplanted MISTRG mouse left untreated after the adoptive transfer of patient-derived immune cells and killed at day 26. Microscopic evaluation: 20X.

### Figure S7 related to Figure 7

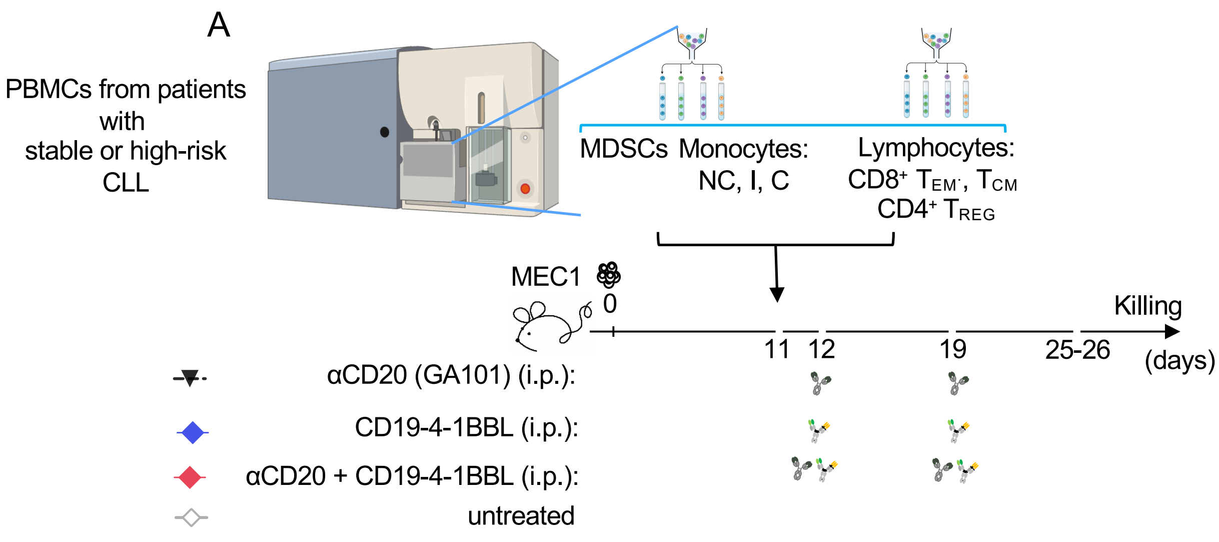


B

high-risk CLL

| Pt ID | CLL Diagnosis (y) | Rai stage | Infection history | WBC | Treat-ment | ZAP70 % # | IGHV MS | FISH |
|---|---|---|---|---|---|---|---|---|
| 446 | 2019 | N/A | N/A | 51 | N | N/A | U | Trisomy12 |
| 110 | 2017 | 0 | N/A | 215.3 | N | - | M | Del(13q) |
| 494 | 2016 | N/A | N/A | 123.2 | N | N/A | U | Del(11q) Del(13q) |

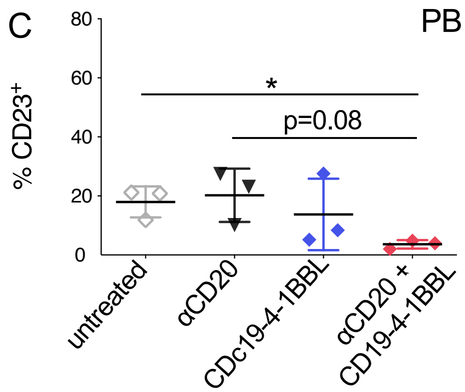

**Fig. S7. Antileukemic effect of CD19-4-1BBL in the MISTRG-based xenograft system**

(A) MISTRG mice transplanted i.v. with MEC1 cells (day 0) received weekly injections of CD19-4-1BBL (1mg/kg, blue rombi) or $\alpha$CD20 (10mg/kg, black triangles) or their combination (red rombi), after the adoptive transfer of patient-derived immune cells (days 11 or day 20, depending on the experiment,) including NC, I, C monocyte subsets, M-MDSCs, $CD4^+$ $T_{REG}$, $CD8^+$ effector $T_{EM}$ and central memory $T_{CM}$ cells separated using fluorescence-activated cell sorting from the PBMCs of patients described in Table B. Mice were euthanized at days 25-26. (B) Clinical features of related patients with high-risk CLL are represented. (C) The mean value of the relative contributions of $CD23^+$ B cells in peripheral blood (PB) is shown in the graph. A statistical analysis was performed using the Student $t$ test $^*P < 0.05$.

### Figure S8 related to Figure 7

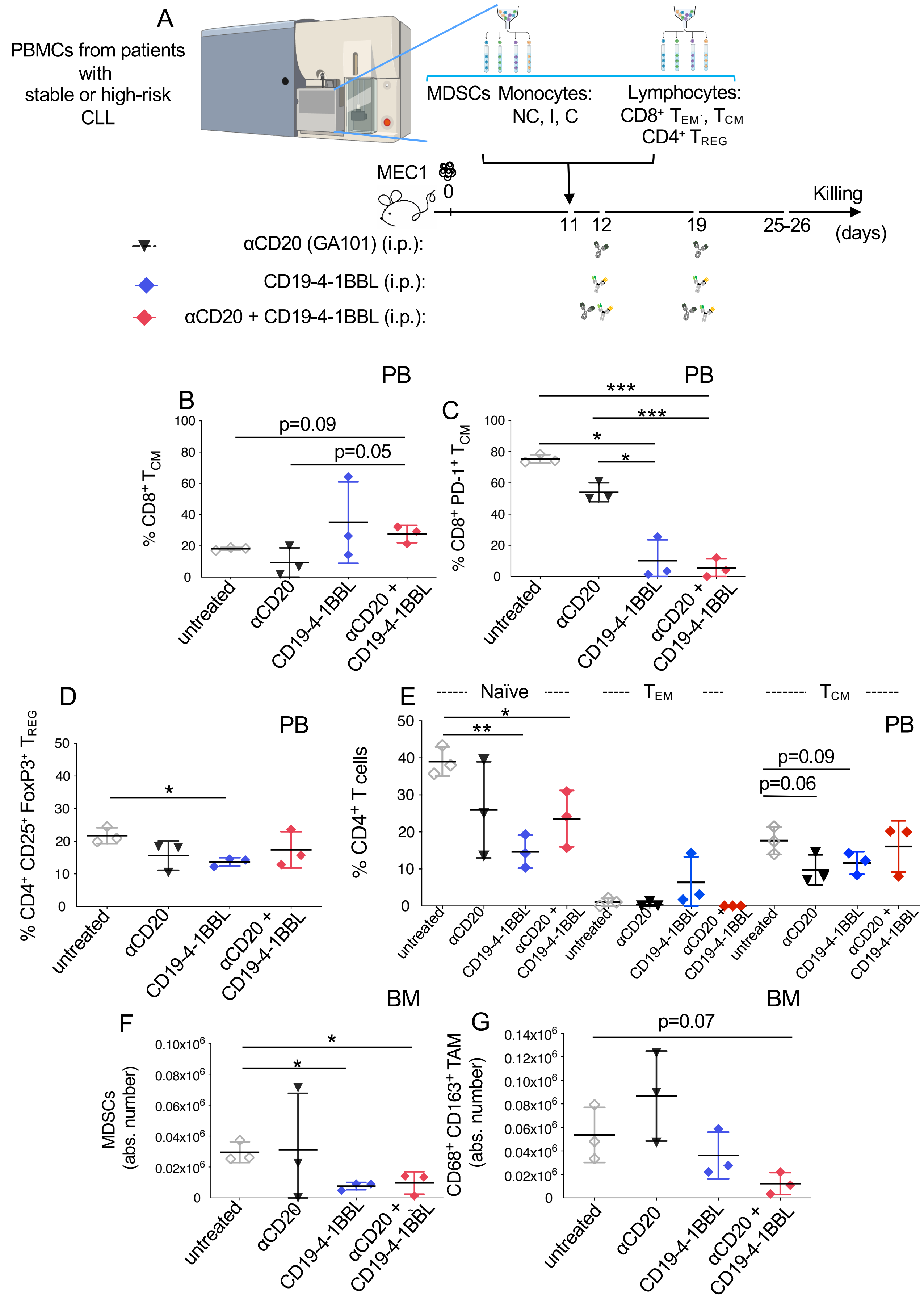

**Fig. S8. The effect of CD19-4-1BBL on patient-derived immune cells of the MISTRG-based xenograft system**

(A) MISTRG mice transplanted i.v. with MEC1 cells (day 0) received weekly injections of CD19-4-1BBL (1mg/kg, blue rombi) or $\alpha$CD20 (10mg/kg, black triangles) or their combination (red rombi), after the adoptive transfer of patient-derived immune cells (days 11 or day 20, depending on the experiment,) including NC, I, C monocyte subsets, M-MDSCs, $CD4^+$ $T_{REG}$, $CD8^+$ effector $T_{EM}$ and central memory $T_{CM}$ cells separated using fluorescence-activated cell sorting from the PBMCs of patients described in Table B. Mice were euthanized at days 25-26.

(B-E) The mean value of the relative contributions of $CD8^+$ $T_{CM}$, $CD8^+$ $PD1^+$ $T_{CM}$, $CD4^+$ $CD25^+$ $FoxP3^+$ $T_{REG}$, $CD4^+$ $CD45RO^-CD45RA^+$ $CD62L^+$ Naïve, $CD4^+$ $CD45RA^-$ $CD45RO^+$ $CD62L^-$ $T_{EM}$, $CD4^+$ $CD45RA^-$ $CD45RO^+$ $CD62L^+$ $T_{CM}$, in peripheral blood (PB) is shown in the graphs. (F-G) The mean values of the absolute numbers of the whole pool of $HLA\text{-}DR^+$ $CD14^{low/neg}$ M-MDSCs and $CD68^+$ $CD163^+$ TAMs in BM are shown in the graphs. A statistical analysis was performed using the Student *t* test $^*P < 0.05$, $^{**}P < 0.01$, $^{***}P < 0.001$.

### Figure S9

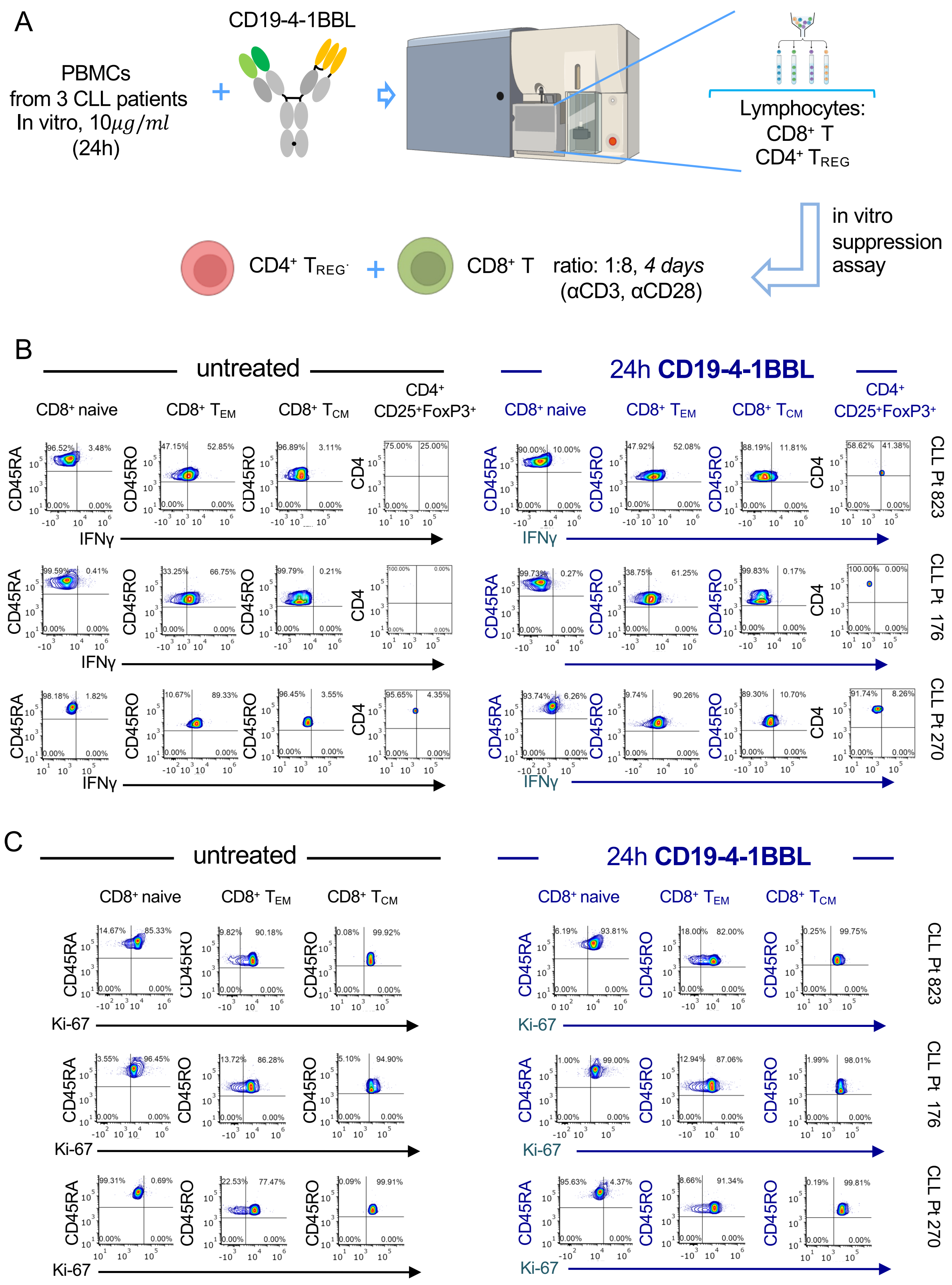

A
CD19-4-1BBL
PBMCs from 3 CLL patients In vitro, 10μg/ml (24h)
Lymphocytes: CD8+ T CD4+ TREG
in vitro suppression assay
CD4+ TREG
CD8+ T
ratio: 1:8, 4 days (αCD3, αCD28)
B
untreated
24h CD19-4-1BBL
CD8+ naive
CD8+ TEM
CD8+ TCM
CD4+ CD25+FoxP3+
CD45RA
CD45RO
CD4
IFNγ
CLL Pt 823
CLL Pt 176
CLL Pt 270
C
untreated
24h CD19-4-1BBL
CD8+ naive
CD8+ TEM
CD8+ TCM
Ki-67

**Fig. S9. The effect of CD19-4-1BBL on the immunosuppressive capacity of patient-derived $CD4^+$ $T_{REG}$ cells on $CD8^+$ T lymphocytes**

(A) Fresh PBMCs from patients with CLL (n = 3, Table S1) were incubated with CD19-4-1BBL (10µ/ml) or left untreated for 24 h. Fluorescent-activated cell sorted (FACS) $CD4^+CD25^+CD127^{low/-}$ regulatory T cells ($T_{REGS}$) and total $CD8^+$ cells were co-cultured in vitro for 4 days at 1:8 ratio ($T_{REG}$ cell numbers: 33,893 for samples 823-176 and 19,741 for sample 270; $CD8^+$ T cell number: 271,144 for samples 823-176 and 157,928 for sample #270). (B) Intracellular spectral flow cytometry analysis of IFN$\gamma$ production by $CD8^+$ $CD45RO^-CD45RA^+CD62L^+$ naive T cells, $CD8^+$ $CD45RA^-CD45RO^+CD62L^-$ $T_{EM}$, $CD8^+$ $CD45RA^-CD45RO^+CD62L^+$ $T_{CM}$ and, $CD4^+$ $CD25^+$ $FoxP3^+$ $T_{REG}$. The percentage of IFN$\gamma$ producing cells is indicated. (C) Spectral flow cytometry analysis of Ki-67 expression on $CD8^+$ $CD45RO^-CD45RA^+CD62L^+$ naive T cells, $CD8^+$ $CD45RA^-CD45RO^+CD62L^-$ $T_{EM}$, $CD8^+$ $CD45RA^-CD45RO^+CD62L^+$ $T_{CM}$ and, $CD4^+$ $CD25^+$ $FoxP3^+$ $T_{REG}$. The percentage of $Ki\text{-}67^+$ cells is indicated.

### Figure S10

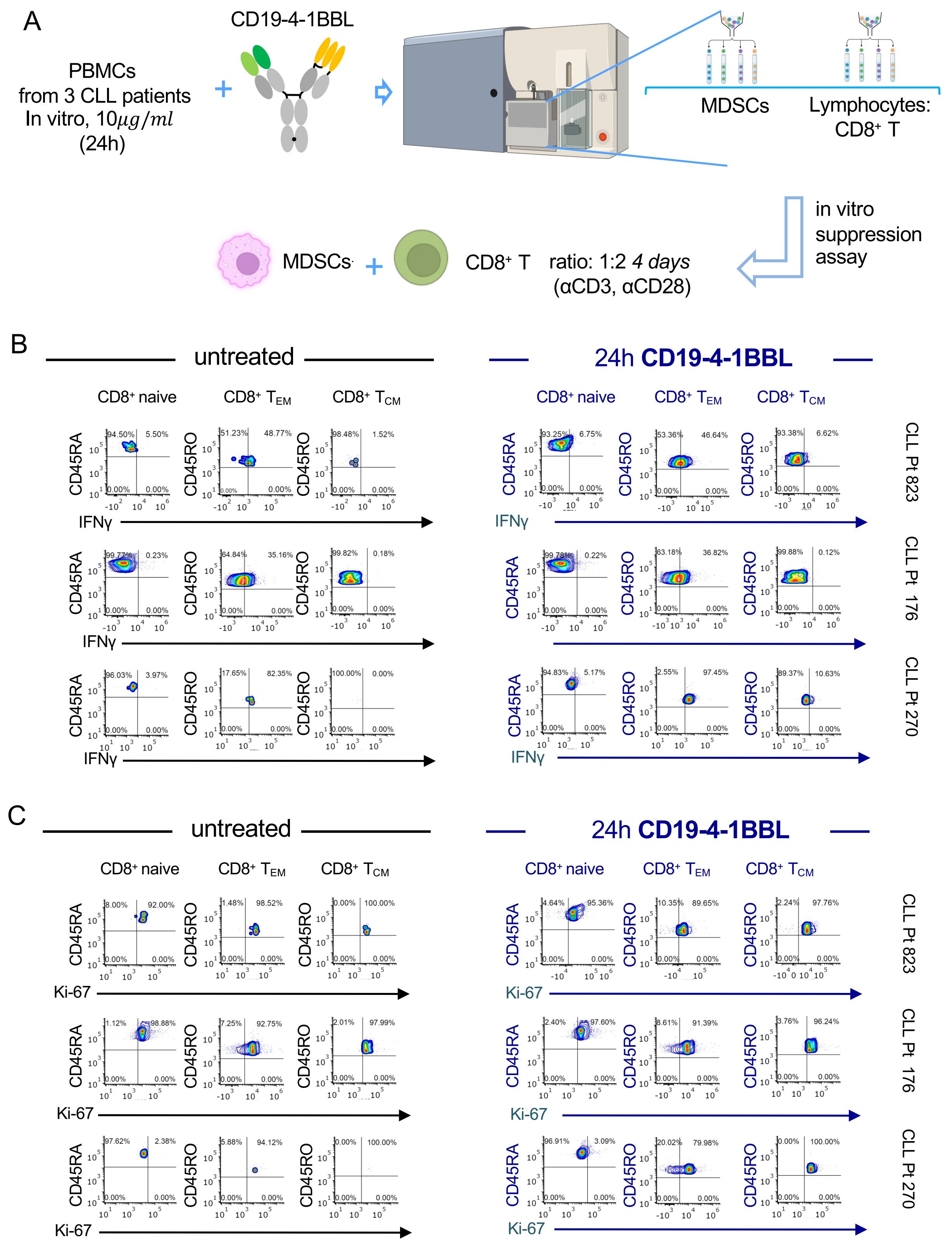
A
PBMCs
from 3 CLL patients
In vitro, 10μg/ml
(24h)
CD19-4-1BBL
MDSCs
Lymphocytes:
CD8+ T
in vitro
suppression
assay
MDSCs
CD8+ T
ratio: 1:2 4 days
(αCD3, αCD28)
B
untreated
24h CD19-4-1BBL
CD8+ naive
CD8+ TEM
CD8+ TCM
CD45RA
CD45RO
IFNγ
CLL Pt 823
CLL Pt 176
CLL Pt 270
C
untreated
24h CD19-4-1BBL
CD8+ naive
CD8+ TEM
CD8+ TCM
CD45RA
CD45RO
Ki-67
CLL Pt 823
CLL Pt 176
CLL Pt 270

**Fig. S10. The effect of CD19-4-1BBL on the immunosuppressive capacity of patient-derived M-MDSCs on $CD8^+$ T lymphocytes**

(A) Fresh PBMCs from patients with CLL (n = 3, Table S1) were incubated with CD19-4-1BBL (10µ/ml) or left untreated for 24 h. Fluorescent-activated cell sorted (FACS) $CD14^+$HLA-$DR^{low/-}$ M-MDSCs and total $CD8^+$ cells were co-cultured in vitro for 4 days at 1:2 ratio (M-MDSC cell numbers: 3249 for samples #823-#176 and 969 for sample #270; $CD8^+$ T cell number: 6498 for samples #823-#176 and 1938 for sample #270). (B) Intracellular spectral flow cytometry analysis of IFN$\gamma$ production by $CD8^+$ $CD45RO^-CD45RA^+CD62L^+$ naive T cells, $CD8^+$ $CD45RA^-CD45RO^+CD62L^-$ $T_{EM}$, $CD8^+$ $CD45RA^-CD45RO^+CD62L^+$ $T_{CM}$. The percentage of IFN$\gamma$ producing cells is indicated. (C) Spectral flow cytometry analysis of Ki-67 expression by $CD8^+$ $CD45RO^-CD45RA^+CD62L^+$ naive T cells, $CD8^+$ $CD45RA^-CD45RO^+CD62L^-$ $T_{EM}$, $CD8^+$ $CD45RA^-CD45RO^+CD62L^+$ $T_{CM}$ and, $CD4^+$ $CD25^+$ $FoxP3^+$ $T_{REG}$. The percentage of Ki-$67^+$ cells is indicated.

#### SUPPLEMENTARY REFERENCES TO TABLE S7

272. Vieyra M, Leisman S, Raedler H, et al. Complement regulates CD4 T-cell help to CD8 T cells required for murine allograft rejection. *Am J Pathol*. 2011;179(2):766-774.
273. West EE, Kemper C. Complement and T Cell Metabolism: Food for Thought. *Immunometabolism*. 2019;1(T Cell Metabolic Reprogramming):e190006.
274. Devi S, Indramohan M, Jäger E, et al. CARD-only proteins regulate in vivo inflammasome responses and ameliorate gout. *Cell Rep*. 2023;42(3):112265.
275. Dumontet E, Osman J, Guillemont-Lambert N, Cros G, Moshous D, Picard C. Recurrent Respiratory Infections Revealing CD8α Deficiency. *J Clin Immunol*. 2015;35(8):692-695.
276. Arroyo-Sánchez D, Luczkowiak J, Delgado R, Cabrera-Marante O, Paz-Artal E. Immune Response Against SARS-CoV-2 Infection and Vaccination in a CD8α-Deficient Patient. *J Clin Immunol*. 2023;43(6):1072-1074.
277. Arenas V, Castaño JL, Domínguez-García JJ, Yáñez L, Pipaón C. A Different View for an Old Disease: NEDDylation and Other Ubiquitin-Like Post-Translational Modifications in Chronic Lymphocytic Leukemia. *Front Oncol*. 2021;11:729550.
278. MacEwan DJ. TNF ligands and receptors--a matter of life and death. *Br J Pharmacol*. 2002;135(4):855-875.
279. Duan H, Dixit VM. RAIDD is a new 'death' adaptor molecule. *Nature*. 1997;385(6611):86-89.
280. Kono M, Komatsuda H, Yamaki H, et al. Immunomodulation via FGFR inhibition augments FGFR1 targeting T-cell based antitumor immunotherapy for head and neck squamous cell carcinoma. *Oncoimmunology*. 2022;11(1):2021619.
281. Palakurthi S, Kuraguchi M, Zacharek SJ, et al. The Combined Effect of FGFR Inhibition and PD-1 Blockade Promotes Tumor-Intrinsic Induction of Antitumor Immunity. *Cancer Immunol Res*. 2019;7(9):1457-1471.
282. Souza-Fonseca-Guimaraes F, Krasnova Y, Putoczki T, et al. Granzyme M has a critical role in providing innate immune protection in ulcerative colitis. *Cell Death Dis*. 2016;7(7):e2302.
283. Krenacs L, Smyth MJ, Bagdi E, et al. The serine protease granzyme M is preferentially expressed in NK-cell, gamma delta T-cell, and intestinal T-cell

lymphomas: evidence of origin from lymphocytes involved in innate immunity. *Blood*. 2003;101(9):3590-3593.

329. Qu Y, Wang X, Bai S, et al. The effects of TNF-α/TNFR2 in regulatory T cells on the microenvironment and progression of gastric cancer. *Int J Cancer*. 2022;150(8):1373-1391.
330. Chang JH, Kim YJ, Han SH, Kang CY. IFN-gamma-STAT1 signal regulates the differentiation of inducible Treg: potential role for ROS-mediated apoptosis. *Eur J Immunol*. 2009;39(5):1241-1251.
331. Ma H, Lu C, Ziegler J, et al. Absence of Stat1 in donor $CD4^+$ T cells promotes the expansion of Tregs and reduces graft-versus-host disease in mice. *J Clin Invest*. 2011;121(7):2554-2569.
332. Gu AD, Zhang S, Wang Y, Xiong H, Curtis TA, Wan YY. A critical role for transcription factor Smad4 in T cell function that is independent of transforming growth factor β receptor signaling. *Immunity*. 2015;42(1):68-79.
333. Proto JD, Doran AC, Gusarova G, et al. Regulatory T Cells Promote Macrophage Efferocytosis during Inflammation Resolution. *Immunity*. 2018;49(4):666-677.e666.
334. Świerzko AS, Michalski M, Sokołowska A, et al. Associations of Ficolins With Hematological Malignancies in Patients Receiving High-Dose Chemotherapy and Autologous Hematopoietic Stem Cell Transplantations. *Front Immunol*. 2019;10:3097.
335. Cibrián D, Sánchez-Madrid F. CD69: from activation marker to metabolic gatekeeper. *Eur J Immunol*. 2017;47(6):946-953.
336. Lu H, Dai X, Li X, Sun Y, Gao Y, Zhang C. Gal-1 regulates dendritic cells-induced Treg/Th17 balance though NF-κB/RelB-IL-27 pathway. *Ann Transl Med*. 2019;7(22):628.
337. Takimoto T, Wakabayashi Y, Sekiya T, et al. Smad2 and Smad3 are redundantly essential for the TGF-beta-mediated regulation of regulatory T plasticity and Th1 development. *J Immunol*. 2010;185(2):842-855.
338. Xue JF, Hua F, Lv Q, et al. DEDD negatively regulates transforming growth factor-beta1 signaling by interacting with Smad3. *FEBS Lett*. 2010;584(14):3028-3034.
339. Churpek JE, Smith-Simmer K. DDX41-Associated Familial Myelodysplastic Syndrome and Acute Myeloid Leukemia. In: Adam MP, Mirzaa GM, Pagon RA, et al., eds. GeneReviews(®). Seattle (WA): University of Washington, Seattle

340. Qin K, Jian D, Xue Y, et al. DDX41 regulates the expression and alternative splicing of genes involved in tumorigenesis and immune response. *Oncol Rep*. 2021;45(3):1213-1225.
341. Gabriel SS, Bon N, Chen J, et al. Distinctive Expression of Bcl-2 Factors in Regulatory T Cells Determines a Pharmacological Target to Induce Immunological Tolerance. *Front Immunol*. 2016;7:73.
342. Tischner D, Gaggl I, Peschel I, et al. Defective cell death signalling along the Bcl-2 regulated apoptosis pathway compromises Treg cell development and limits their functionality in mice. *J Autoimmun*. 2012;38(1):
